# An explanatory benchmark of spatial domain detection reveals key drivers of method performance

**DOI:** 10.64898/2026.03.12.710462

**Authors:** Alice Descoeudres, Tomislav Prusina, Noah Schmidt, Van Hoan Do, Simon Mages, Johanna Klughammer, Domagoj Matijevic, Stefan Canzar

## Abstract

The spatial organization of cells within tissues is critical for understanding biological function and disease, and spatial transcriptomics enables genome-wide mapping of this organization. Numerous computational methods aim to identify spatial domains, yet their performance is often evaluated on limited datasets, leading to conflicting conclusions. Here, we present an explanatory benchmark of 26 spatial domain detection methods across 63 tissue sections from six spatial transcriptomics technologies, supplemented by over 1,000 semi-synthetic datasets that systematically vary resolution, gene panel size, and tissue architecture. By jointly analyzing real and semi-synthetic data across this broad parameter space, our benchmark uncovers systematic performance differences and sources of variability that are obscured in standard evaluations. Although most spatial methods outperform non-spatial baselines, their performance depends strongly on data resolution and cellular heterogeneity. To enable systematic analysis beyond individual methods, we introduce a modular, plug-and-play benchmarking framework that facilitates method refinement and component exchange. Using this framework, an ablation study of neural network-based approaches shows that the choice of preprocessing and clustering often has a larger impact on performance than architectural novelty alone. Together, these results provide a principled foundation for informed method selection and offer guidance for the development of robust and scalable spatial domain detection tools as spatial transcriptomics technologies continue to advance.

## Introduction

Spatial transcriptomics (ST) approaches have enabled researchers to profile gene expression within the original tissue context. Techniques range from targeted, single-molecule resolution fluorescence in situ hybridization (FISH)-based approaches [1–3] to grid-based full-transcriptome profiling methods utilizing next generation sequencing. Increasingly, the field is shifting toward a best-of-both-worlds scenario, with highly multiplexed or full-transcriptome technologies at near single-cell resolution [4–6]. The wealth of data created with these technologies is expected to bring about new insights in many fields, such as cancer research [7], development [8], and neuroscience [9].

A fundamental step in the analysis of ST data is the partitioning of tissue into spatial domains. Domain definitions provide a structured framework to characterize spatial organization. They enable the derivation of domain-specific gene expression signatures [10] and cell type compositions [11], support the mapping of tissue architecture [12], and allow comparative analyses across conditions, technologies, and species. Spatial domains also serve as anchor units for integrative analyses with histology, proteomics, or single-cell RNA sequencing, which makes domain detection a cornerstone of most downstream ST workflows. This has motivated the development of numerous computational approaches for spatial domain detection.

The evaluation of new computational methods is limited and biased, a problem that has been noted repeatedly in the literature, including in previous benchmarking studies across a range of data analytical tasks [13–16]. Reported performance results in original method publications are typically based on narrow datasets and specific evaluation metrics, which can favor the newly introduced method and make comparisons inconsistent across studies. For spatial domain detection, these issues are particularly visible because most methods have been evaluated on the same 10x Genomics Visium dataset from the layered human dorsolateral prefrontal cortex published by Maynard *et al.* [17]. Collectively, the performance claims in existing benchmarks are contradictory. These inconsistencies become apparent when visualized as cycles in a directed graph representing reported pairwise performance superiority (Supplementary Figs. 1 and 2).

Several benchmarking studies have attempted to address this problem in the context of spatial domain detection [18–21]. While these efforts considered a broader range of datasets and evaluation metrics than individual tool publications, they failed to resolve the contradictions and, in some cases, reported conflicting conclusions. These issues arise from the exclusive reliance on real datasets for benchmarking. As a result, existing benchmarks remain largely descriptive and offer limited insight into why methods succeed or fail under different conditions. Ground-truth domain annotations are typically manually curated and subject to considerable uncertainty, which affects performance estimates. Furthermore, any finite set of real datasets covers only a small part of the space of experimental and biological parameters that influence method performance. This incomplete coverage prevents a comprehensive understanding of how methods behave under different technological or tissue conditions. To address these limitations, our benchmark uses real datasets to identify patterns in method performance and then systematically evaluates these patterns using semi-synthetic data. The simulations allow independent variation of technological parameters such as resolution, sparsity, and gene panel size, as well as tissue properties such as domain size and shape, cell type distinctness, and compositional heterogeneity between domains. This systematic exploration of parameter space reveals patterns that cannot be observed from real data alone.

We also analyze the impact of internal stochastic processes implemented by many methods, such as random weight initialization or data augmentation during contrastive learning. These components can cause the same method to produce different outputs across different trials on the same dataset, contributing to the contradictory results reported across publications. Because many implementations do not expose controls for these random elements, their effect on performance has largely been overlooked in previous benchmarks. To address this, we introduce a controlled perturbation strategy by permuting the input cell order without altering the underlying data to quantify the inherent stochasticity of each method and assess its impact on performance stability.

Beyond benchmarking, we provide a modular, open-source software framework that supports consensus analysis and facilitates method development. Using this framework, we show that consensus domains derived from all included methods consistently outperform individual methods on Visium datasets, while performing competitively on high-resolution data. Finally, an ablation study disentangles the contributions of neural network architecture, preprocessing, and clustering to overall performance. The resulting plug-and-play framework enables method developers to refine individual components, such as neural network loss functions, while integrating them seamlessly with established preprocessing and clustering strategies.

By combining systematic simulation with real data benchmarking, we uncover hidden sources of variability and performance trade-offs that remain invisible in conventional evaluations. Our results offer actionable guidance for users selecting methods and for developers designing the next generation of spatial analysis tools.

## Results

### Benchmark overview

We benchmarked 26 methods for spatial domain identification, which we grouped into four broad categories: clustering-based, neural network-based, statistical model-based, and image segmentation-based approaches (Fig. 1a). Clustering-based methods construct a joint representation of gene expression and spatial coordinates and infer domains using standard clustering algorithms. Most neural network-based methods employ graph convolutional networks (GCNs) to learn low-dimensional embeddings that capture both spatial and transcriptional structure. Statistical model-based methods incorporate spatial information directly into probabilistic models of gene expression, using hidden Markov random fields (HMRF) or nonnegative matrix factorization (NMF). Vesalius encodes gene expression as RGB values and applies a classical image segmentation pipeline. A conceptual overview of the neural-network-based methods is provided in Supplementary Note 1, and method-by-method descriptions of all approaches are given in the Methods. As baselines, we included the non-spatial Leiden algorithm as implemented in scanpy [22] and Seurat [23]. Additionally, a simple spatial smoothing strategy applied to these non-spatial clustering outputs serves as a naïvely spatially-aware baseline (Methods).

**Figure 1:**
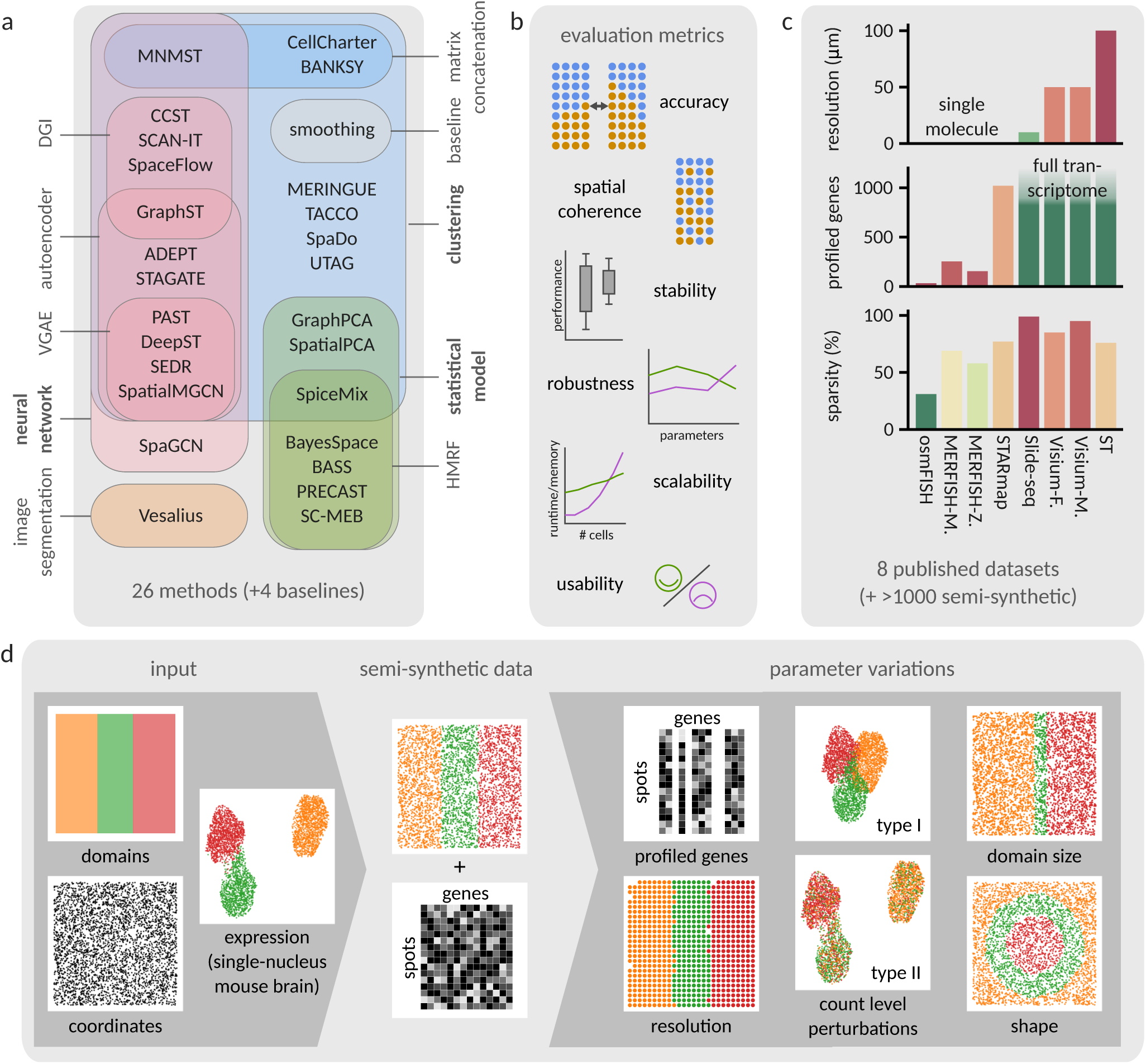
Overview of the benchmark. **a**, Categorization of spatial domain detection methods included in the benchmark. **b**, Metrics used for method evaluation. **c**, Overview of published datasets in terms of resolution, number of profiled genes, and sparsity of the resulting count matrix. **d**, Schematic of the generation of semi-synthetic data. Domain shapes and cell coordinates generated *in silico* are combined with gene expression from a publicly available single-nucleus RNA-seq dataset of the mouse brain. The resulting semi-synthetic dataset can be further modified, for example with respect to technological parameters, count level perturbations, or tissue architecture.

We evaluated the performance of all methods using a diverse set of metrics (Fig. 1b and Methods). As a first step, methods ran on several publicly available datasets spanning different tissues and generated using diverse technologies, including both sequencing-based and image-based platforms (Fig. 1c, Methods, and Supplementary Note 2). All methods were executed using the developer-recommended, technology-specific parameter settings (Methods). Accuracy of methods’ output was assessed using the Adjusted Rand Index (ARI; see Methods) against expert manual annotations serving as ground truth. The datasets contain between 266 (Spatial Transcriptomics) and over 5,500 (MERFISH) cells or spots, and vary in resolution, capture sensitivity, and size of the profiled gene panel, ranging from 33 genes in osmFISH to full-transcriptome coverage in sequencing-based datasets, reflecting the technological heterogeneity across spatial transcriptomics platforms. Beyond method accuracy, we develop an approach to evaluate method stability without relying on random seed initialization. This enables stability assessment even for methods that do not expose a seed or control over their internal randomness.

Based on method performance on publicly available datasets, we identified recurring patterns and performance trends associated with specific data characteristics. To systematically examine these patterns, we developed a highly flexible pipeline for generating semi-synthetic spatial transcriptomics datasets (Fig. 1d). The pipeline generates cell coordinates with ground-truth domain annotations across a range of domain shapes and tissue conformations. Counts are assigned from cell type profiles previously identified in a single-nucleus RNA-seq dataset of the mouse brain [24]. The semi-synthetic base data can be modified by different types of noise, changing cell type compositions, downsampling genes, or aggregating cells into spots. To avoid shape-specific biases, we simulated multiple conformations (Methods) and report results of accuracy averaged across shapes, with potential shape effects examined separately. The full description of the pipeline is provided in Methods.

Importantly, the methods are also evaluated based on scalability in terms of runtime and memory usage, and usability. Scalability is evaluated using synthetic datasets of various sizes, generated by SRTsim [25]. For usability evaluation, we adapt a weighted checklist from [26] and score methods according to the quality of their code and documentation, ease of installation, and continued maintenance.

Using our comprehensive Snakemake pipeline, we investigate two analysis approaches beyond individual method output. We perform a modularization-based ablation analysis of six neural network-based methods to investigate whether individual components have a disproportionate effect on method performance. Additionally, we show that combining different methods through a consensus approach increases robustness across technologies and can improve performance by integrating complementary information captured by different algorithms.

### Not all spatial methods outperform non-spatial baselines

To assess the added value of spatial modeling, we first compare spatial methods against non-spatial baselines. The clustering accuracy achieved by non-spatial methods reflects how well expert-annotated spatial domains align with clusters defined purely by gene expression. Across datasets, every spatial method outperforms the non-spatial baselines at least once, although in some cases the improvements are only marginal (Fig. 2a).

**Figure 2:**
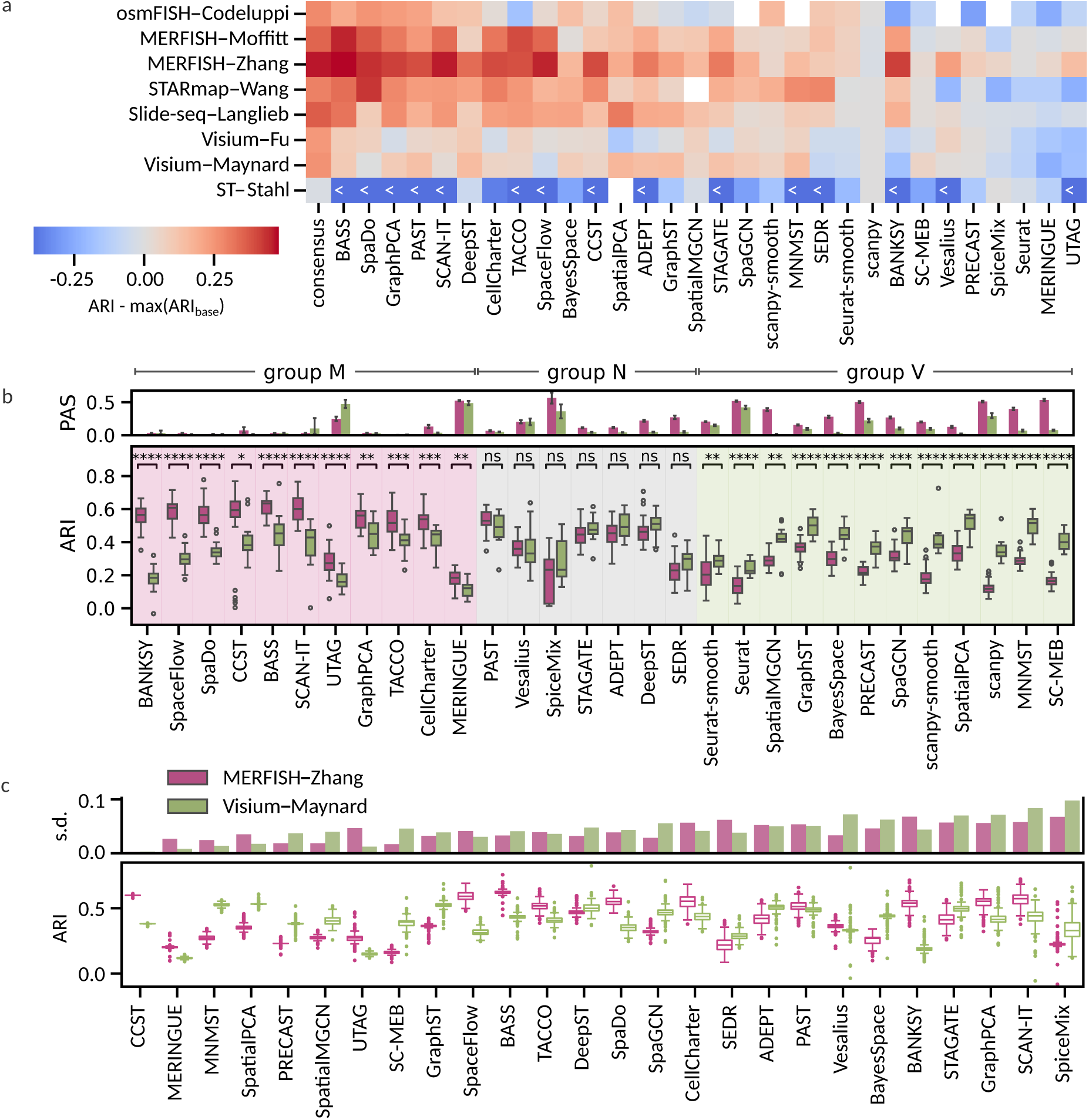
Method performance on real datasets. **a**, Difference in Adjusted Rand indices (ARI) between each spatial method and the best-performing non-spatial baseline (scanpy or Seurat). Datasets are ordered by spatial resolution, and methods are ordered by their mean standardized ARI across datasets. The color scale is truncated at –0.4 to highlight the most relevant performance differences, with the truncation effect marked with “*<*” in the affected cells. **b**, ARI values of all methods on the Visium-Maynard and MERFISH-Zhang datasets. Methods are ordered by the difference in mean ARI between the two technologies and grouped by statistical significance. Statistical significance was assessed using two-sided Mann-Whitney U tests. Corresponding Percentage of Abnormal Spots (PAS) values for each method on the same datasets are shown in the top panel. **c**, Stochastic variability in domain detection accuracy. For each method, ARI values are shown across 12 random trials on each of the 12 Visium-Maynard and 12 MERFISH-Zhang samples. Methods are ordered by their mean standard deviation in ARI across the two datasets (top bar plot). ARI values are normalized to correct for inter-sample variability (Methods). Box plots: boxes show quartiles, whiskers extend to points within 1.5 interquartile lengths of upper and lower quartiles.

Nearly one-third of methods (9 of 26) perform worse overall than a simple spatial smoothing applied to the scanpy clustering output, as measured by mean standardized ARI. The smoothing improves scanpy’s clustering accuracy by up to 0.1 ARI on the STARmap and osmFISH datasets. Interestingly, on the ST-Stahl dataset, smoothing reduces baseline performance by as much as 0.2 ARI, whereas non-spatial baselines achieve ARIs above 0.95. Since all spatial methods except SpiceMix [27] are outperformed by the baselines on this dataset, we exclude it from further analysis. The extent of improvement provided by spatial methods varies substantially across datasets. In the low-resolution Visium datasets, performance gains over non-spatial methods are modest (maximum ARI increase of 0.16), whereas in higher-resolution datasets, spatial methods potentially yield much larger improvements (ARI increases of 0.48 on MERFISH and 0.32 on Slide-seq). In addition, aggregating the outputs of all methods using a consensus strategy (Methods) yields accuracies that consistently exceed those of any individual method. The consensus performs competitively on high-resolution datasets and shows particularly large gains over the baselines on the Visium datasets.

Beyond resolution, other technological factors also appear to influence the effectiveness of spatial domain detection methods. One such factor is detection sensitivity: Slide-seq, which exhibits high count sparsity, shows consistently poor absolute performance across nearly all methods (Supplementary Fig. 3). Another important variable is the size of the gene panel used in targeted approaches. For instance, osmFISH includes only 33 genes in its panel, and across methods, it shows smaller gains over non-spatial baselines compared to other high-resolution technologies. These observations suggest that resolution, detection sensitivity, and gene panel size can substantially limit the benefits of spatial modeling, which we explore more systematically using semi-synthetic data.

### Spatial coherence and performance correlate on high-resolution data

To further investigate the observed performance differences between technologies, we next focused on Visium and MERFISH, two widely adopted spatial transcriptomics technologies that represent opposite extremes in both spatial resolution and number of profiled genes. In fact, the relevance of spatial information varied substantially between the two (see improvement over non-spatial methods in Fig. 2a, and performance of non-spatial methods in Supplementary Fig. 3). For this analysis, we selected the Visium-Maynard and MERFISH-Zhang datasets, as both contain more than 10 samples and exhibit comparable laminar tissue structure in the human and mouse brain, respectively. Figure 2b shows that the majority of methods (groups M and V) exhibit highly significant performance differences between the two datasets. To help explain these differences, we assessed spatial coherence, quantified by the Percentage of Abnormal Spots (PAS; see Methods). Across methods, PAS values are significantly higher on the MERFISH data, indicating that the inferred domains tend to be less spatially coherent than on Visium (Supplementary Fig. 4). Notably, most methods that perform significantly better on MERFISH than on Visium (group M) display high spatial coherence, evidenced by low PAS, in both datasets. The two exceptions UTAG [28] and MERINGUE [29] generally show weaker performance. In contrast, methods that perform significantly better on the lower-resolution Visium dataset (group V) exhibit higher PAS overall, and especially on MERFISH. This pattern suggests that the accuracy of a given clustering is closely tied to its spatial coherence in high-resolution datasets, as supported by a strong median Spearman correlation of *−*0.85 between ARI and PAS in high-resolution data (Supplementary Fig. 5).

Methods from group V generally maintain high spatial coherence on the lower-resolution Visium data, even if their coherence is reduced in high-resolution settings. However, the converse does not hold: low PAS values do not necessarily imply high clustering accuracy. This asymmetry is reflected by a weaker median Spearman correlation of *−*0.31 between ARI and PAS (Fig. 2b, Supplementary Fig. 5). These observations suggest that differences in intrinsic spatial smoothness between technologies interact with method design: methods that impose stronger spatial coherence perform best on MERFISH, where expression patterns are less spatially smooth, whereas methods that rely more heavily on transcriptional similarity perform relatively better on Visium. We examine this observation further using our semi-synthetic simulation study.

A comparison of methods based solely on quantitative metrics such as ARI and PAS cannot fully capture the factors influencing performance, such as methods consistently failing to detect specific domains. For instance, in sample 151675 of the Visium-Maynard dataset, none of the top-performing methods correctly identify the L4 domain of the dorsolateral prefrontal cortex (Supplementary Fig. 6). Below we systematically investigate different factors that may impact the ease of detection of specific domains, such as transcriptional distinctness, domain shape, and tissue layout, using semi-synthetic data.

### Stochastic effects on spatial domain identification

Many domain detection methods make random choices, such as random weight initialization or data augmentation during contrastive learning, which can influence the outcome of the analysis. Changing the random seed modifies the sequence of pseudo-random numbers used, potentially leading the algorithm along a different computational path and resulting in variability in the output. However, as noted in previous benchmarks [18], several methods internally fix the random seed, leading to identical choices for the same input and thus concealing the algorithm’s inherent stochasticity. To expose this hidden variability, we permute the order of cells in the input count matrix while preserving their expression values and spatial coordinates (Methods). This procedure forces the algorithm to make different random choices, even when using the same seed. We perform 12 such input permutations for 12 slides each from the Visium-Maynard and MERFISH-Zhang datasets, and quantify the resulting variability in clustering accuracy as shown in Fig. 2c.

CCST [30] exhibits minimal variability across runs on both datasets, with a mean standard deviation in ARI below 0.003. Among the methods performing better on Visium (group V, see Fig. 2b), MNMST [31] and SpatialPCA [32] show very high stability, notably on Visium. From group M, BASS [33], SpaceFlow [34], TACCO [35], DeepST [36], and SpaDo [37] are relatively stable, each with a mean standard deviation below 0.042. In contrast, although STAGATE [38], GraphPCA [39], and SCAN-IT [40] perform competitively on both datasets, they are generally less stable (mean s.d.*>* 0.063). SpiceMix stands out with particularly unstable performance, exhibiting a mean standard deviation of 0.088. Interestingly, stochastic variability across methods seems to be driven less by algorithmic design and more by pipeline decisions such as feature selection, regularization, and postprocessing (Discussion).

### Method robustness to key technological parameters

Using our semi-synthetic data generation pipeline (Methods), we first examined how technological characteristics influence method performance. In particular, we assessed the effect of (i) resolution, by aggregating individual cells into grid-defined spots, (ii) panel size, by downsampling the number of measured genes, and (iii) sparsity, by increasing the initial matrix sparsity of 85% up to 99% through randomly setting counts to zero. The synthetic data cover a wide range of resolutions (Supplementary Fig. 7): grid length 10 corresponds to very low-resolution, ST-like data, whereas grid length 5 yields roughly 9 cells per spot, closely matching the resolution of Visium [17]. A grid length of 0.5 yields spots that closely match the original single-cell locations.

For most methods, performance declines as spatial resolution decreases (representative methods in Fig. 3a; all methods in Supplementary Fig. 8). Some methods, such as BASS, TACCO, STAGATE, and SpiceMix, are affected only at larger spot sizes (grid length *>* 5). Many methods from group M (compare Fig. 2b), such as BANKSY [41], SpaceFlow, and SCAN-IT, show strong and early performance degradation with cell aggregation. In contrast, baseline methods, SpatialMGCN [42], and SpaGCN [43], all of which perform better on Visium than on MERFISH, can even see performance gains when resolution is slightly reduced.

**Figure 3:**
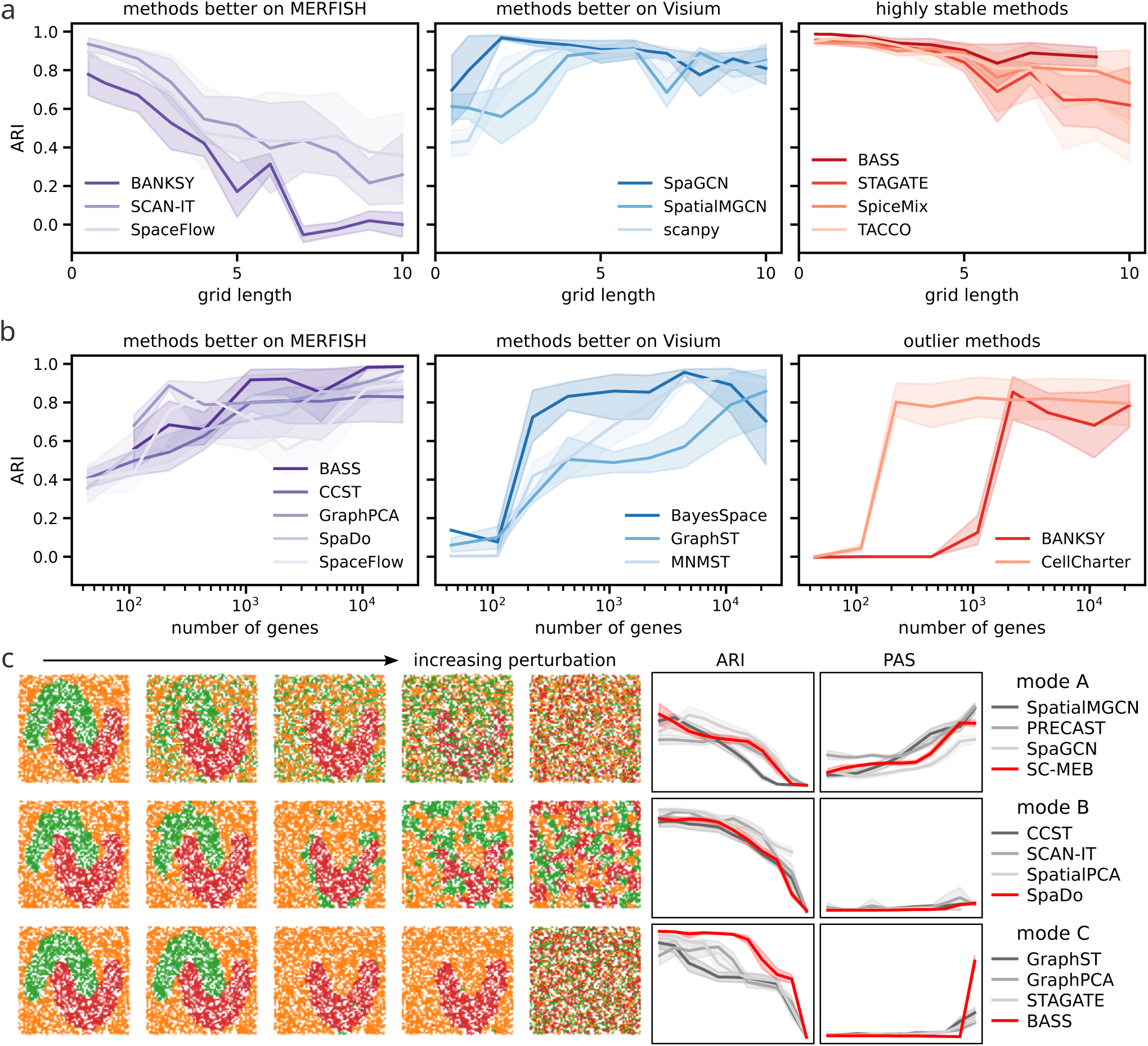
**a**, Performance as a function of spatial resolution (grid side length) for selected methods from different groups. **b**, Performance as a function of the number of profiled genes for selected methods. The *x*-axis is shown on a logarithmic scale. Across a and b, variance reflects differences in tissue layout. **c**, Failure archetypes A, B, and C under type I (ambient RNA) perturbation. ARI and PAS are shown for four representative methods illustrating each archetype, with example domain maps displayed for the methods highlighted in red (representative perturbation levels shown per mode). In the line plots, variance reflects differences in tissue layout (domain shapes) and five random seeds used for count assignment (see Methods). In all domain maps, orange, green, and red spots correspond to domains defined by cell types 1, 2, and 3, respectively (Table 2). Across all panels, solid line shows mean, spread shows 95% confidence interval.

The strong negative ARI-PAS correlation observed on high-resolution real datasets is corroborated by the semi-synthetic results (Supplementary Fig. 9): At no aggregation or small grid lengths, the Spearman correlation between ARI and PAS is –0.55 (median on real high-resolution data: –0.83), whereas at very low resolution the correlation disappears (–0.05).

As for the dependence on gene panel size (representative methods in Fig. 3b; all methods in Supplementary Fig. 8), all competitive methods show decreasing performance as the number of profiled genes shrinks. Most of the methods from group V, such as GraphST [44], BayesSpace [45], and MNMST, approach ARI values near zero at the smallest panel sizes. In contrast, methods from group M, including SpaceFlow, SpaDo, BASS, and CCST, exhibit a more moderate decline. Two notable exceptions are BANKSY and CellCharter [46], whose performance drops sharply at around 1000 and 100 genes, respectively. Both of these methods notably perform worse on the 33-gene osmFISH dataset than on MERFISH with its comparatively larger gene panels (Fig. 2a). Several methods, including BASS, GraphPCA and SpaGCN, did not produce outputs on the smallest gene panels, while SCAN-IT, TACCO, and SpatialMGCN already failed at panel sizes under 1,000 (or 10,000, in the case of SpatialMGCN) genes. Finally, many methods are highly sensitive to extreme count matrix sparsity, including TACCO, PAST [47], STAGATE, SEDR [48], SC-MEB [49], and SpiceMix, whereas others, including BASS, SpatialPCA, GraphPCA, SpaceFlow, MNMST, and SpaDo, are notably more robust (Supplementary Fig. 10). Most of these sparsity-resistant methods perform better on Slide-seq data than would be expected from their performance on other datasets (Fig. 2a).

### Impact of cell type distinctness and compositional heterogeneity on method robustness

Beyond technologically determined data characteristics, we also examined the impact of two types of biological perturbations that may weaken the clustering signal (type I and type II; see Methods for details). In type I, the distinctness of the cell types forming the domains was reduced by adding a common background signal to all cells or spots. This can be interpreted biologically as a continuum of transcriptionally similar cell types, or technically as artifacts such as ambient RNA contamination. In type II, a fraction of cells across the tissue section was assigned to a different cell type, representing infiltrating immune cells or migrating cells during tissue development. Alternatively, this perturbation can be viewed as varying levels of compositional heterogeneity. In both models, 100% perturbation corresponds to the absence of any clustering signal.

Examining the relationship between ARI and PAS under type I perturbation, we identified three archetypal modes of failure (Fig. 3c, with the full set of methods shown in Supplementary Fig. 11). In mode A, a decrease in ARI is mirrored by an increase in PAS, indicating that methods such as SpaGCN, SC-MEB, PRECAST [50], and SpatialMGCN produce increasingly noisy domain maps with blurred boundaries. In mode B, ARI decreases monotonically while PAS remains low, reflecting the appearance of many small disconnected patches (SpatialPCA, SCAN-IT, SpaDo, CCST). In mode C, methods exhibit a sharp drop in ARI caused by a sudden wholesale “flip” of one domain into the label of a neighboring domain. After this flip, ARI enters a plateau around 0.5-0.6, reflecting that the same collapsed configuration is maintained under further perturbation (BASS, STAGATE, GraphPCA, and GraphST). These three modes can be interpreted as different degrees of spatial coherence imposed by the methods: mode A reflects minimal smoothing and high sensitivity to small transcriptional differences, mode B reflects moderate smoothing that fragments domains into small patches, and mode C reflects excessive smoothing that merges or flips whole domains. The domains that collapse in mode C are defined by transcriptionally highly similar cell types 1 and 2 (see Supplementary Fig. 12 and Methods), which are already indistinguishable by baseline methods at zero perturbation (Supplementary Fig. 13). While many methods display mixtures of these behaviors, BASS, ADEPT [51], and TACCO distinguish themselves by maintaining nearly constant median ARIs even at perturbation levels up to 60%. More generally, many methods retain their accuracy under low proportions of type I perturbations.

When applied to tissues with increasing levels of type II perturbations, however, most methods show an immediate decline in accuracy accompanied by rising PAS (Supplementary Fig. 14). PAS values increase to significantly higher levels than under type I perturbations (Supplementary Fig. 15), potentially indicating a strong effect of transcriptional heterogeneity on the visual coherence of domains. Notably, BASS, TACCO, and SpaDo remain more robust than other methods, maintaining performance even with 50-70% infiltrating cell content.

### Impact of domain size and shape on method performance

To assess how domain size influences method performance, we generated semi-synthetic datasets consisting of three laminar layers (Fig. 4a). Starting from a narrow middle layer, we progressively shifted the boundary between layers 2 and 3 toward the rightmost edge of the slice, thereby increasing (decreasing) the thickness of the middle (right) domain. We evaluated detection accuracy for the middle and right domains using the F-score, defined as the geometric mean of recall and precision (Methods). Each experiment was performed under two cell type arrangements: in arrangement A, transcriptionally similar cell types were assigned to the middle domain and its left neighbor, whereas in arrangement B, similarity was assigned to the two outer domains, leaving the middle domain transcriptionally distinct.

**Figure 4:**
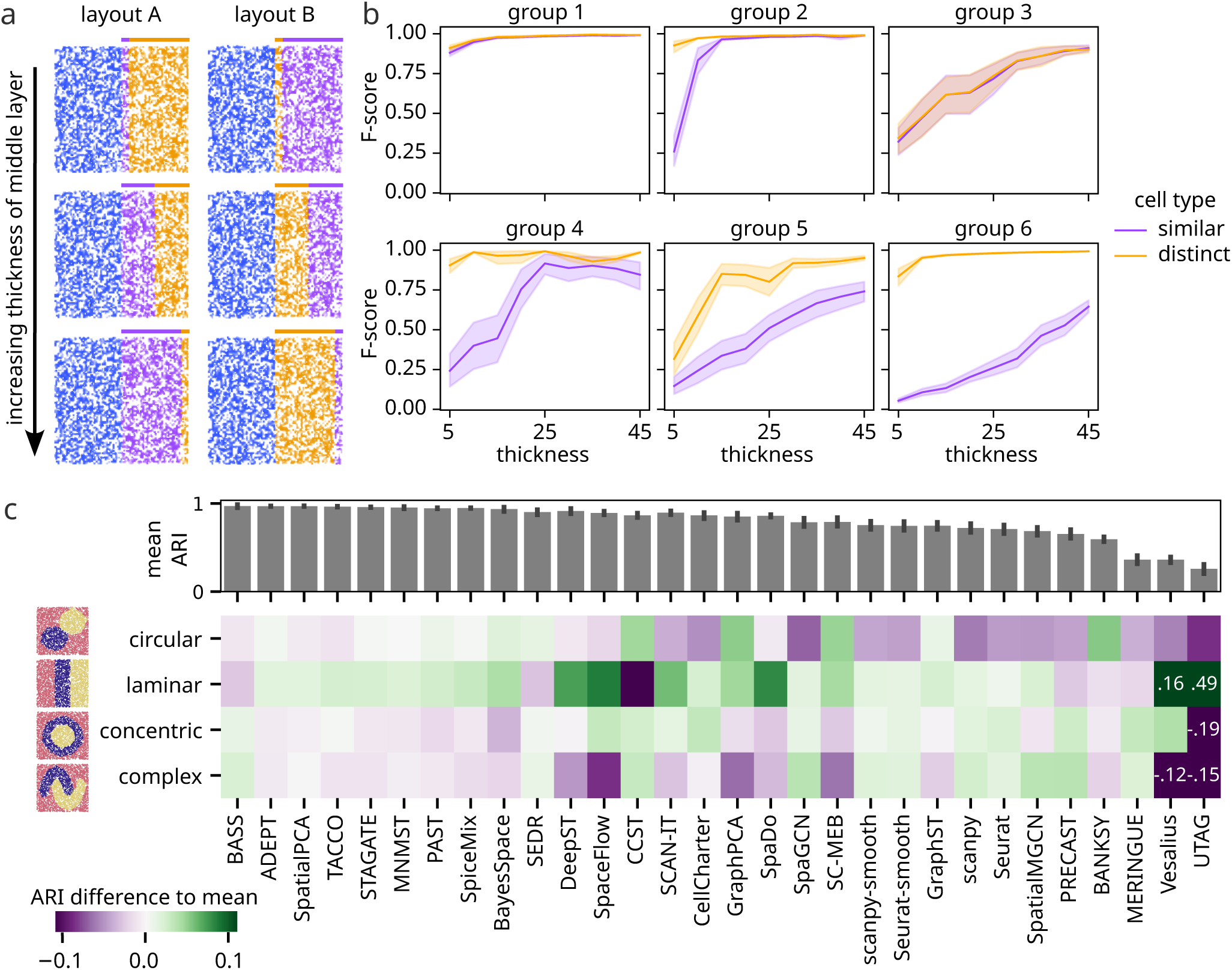
**a**, Representative semi-synthetic samples used to examine the effects of domain thickness and transcriptional similarity. A fixed reference domain (blue, left) is combined with two domains whose transcriptional similarity differs between layouts. In layout A the middle domain is similar (purple) and the right domain distinct (yellow), whereas in layout B the order is reversed. The thickness of the middle domain increases from top to bottom. **b**, Detection accuracy (F-score) for the purple (transcriptionally similar to the reference) and yellow (transcriptionally distinct to the reference) domains across both layouts as a function of domain thickness. Variance reflects five ran-dom seeds used for count assignment (see Methods). Methods are grouped according to their performance patterns. Group 1: BASS, ADEPT, TACCO, SpiceMix, SpaceFlow, SCAN-IT; group 2: DeepST, SpatialPCA, STAGATE, PAST, MNMST; group 3: SpaDo, CellCharter, UTAG, Vesalius; group 4: BayesSpace, SC-MEB, GraphST, SEDR; group 5: GraphPCA, SpaGCN, SpatialMGCN, BANKSY, MERINGUE; group 6: baselines, PRECAST, CCST. Solid line shows mean, spread shows 95% confidence interval. **c**, ARI scores for all methods across different tissue layouts, shown relative to each method’s mean ARI across all shapes. The color scale is truncated at *±*0.11; values exceeding this range are annotated in the heatmap.

Domain thickness had a strong impact on detection accuracy for nearly all methods (Supplementary Fig. 16). In most cases, this effect was modulated by transcriptional distinctness, giving rise to six characteristic behavior patterns (method groups 1-6, Fig. 4b). Methods in group 1 detect the middle domain robustly across both arrangements, with only a minor decline in accuracy at the smallest thickness. Group 2 methods reliably identified domains defined by highly distinct cell types even when they were thin, but struggled when the same domain was transcriptionally similar to its neighbor. Group 4 methods already showed deteriorating performance at moderate thickness in this more challenging scenario, whereas group 5 methods exhibited strong size dependence even when domains were transcriptionally distinct. In contrast, group 3 methods were sensitive to domain size but largely insensitive to transcriptional similarity. Finally, group 6 methods displayed baseline-like behavior, with performance driven primarily by transcriptional distinctness rather than spatial extent. These effects were not restricted to laminar structures. When simulating circular domains of varying diameter, we observed higher proportions of misassigned spots for smaller domains (confusion, defined in Methods, see Supplementary Fig. 17).

Semi-synthetic datasets with different tissue configurations (circular, laminar, concentric, and a complex inter-locking arrangement, as shown in Fig. 4c) revealed that most methods perform best on laminar arrangements. In particular, DeepST, SpaceFlow, SpaDo, and Vesalius showed a clear preference for layered tissues. UTAG performed competitively on layered configurations but failed on all other shapes. Consistent with these findings, several of the best-performing methods on real data, including methods listed above, achieved higher accuracy on the MERFISH-Zhang dataset, which consists of laminar brain layers, than on the structurally more complex MERFISH-Moffitt dataset (Supplementary Figs. 3 and 18).

A consolidated summary of these simulation results and their correspondence to observations from real data is provided in Supplementary Table 1.

### Ablation study of neural network-based methods

All neural network-based methods except SpaGCN compute spatial domains by clustering a low-dimensional embedding of the data (Methods). Specifically, these method architectures can be represented as a series of four steps (see conceptual overview in Supplementary Note 1): After preprocessing the expression data, the spatial adjacency of spots is typically represented by a graph. The neural network computes an embedding of expression and spatial information, which is finally clustered using standard algorithms such as k-means or model-based clustering. Here, we designed experiments to investigate which algorithmic components are crucial for the performance of a method, and whether individual components can improve the quality of solutions if used within other methods. To this end, we selected six methods, two autoencoder-based methods and four Deep-Graph-Infomax-based methods, whose implementation allowed us to split the four components into four independent modules (Methods). We then swapped the implementation of each component with the corresponding module of one of the five other methods while keeping the remaining three components fixed and ran the resulting combination of modules on the 12 Visium-Maynard samples (schematic in Fig. 5a). Fig. 5b shows the maximum absolute change in accuracy when replacing one of the four components (columns) of a given method (row) by a different module. With the exception of GraphST, the choice of adjacency representation and neural network architecture had very little impact on method performance. GraphST was the only method that consistently performed worse when replacing its neural network by a different design or learning objective and it dropped most when swapping it for SEDR’s autoencoder architecture (Supplementary Fig. 19). The other way around, SEDR benefited slightly when using DGI-based neural networks as implemented in SpaceFlow or SCAN-IT instead of its own autoencoder network. In contrast, performance of STAGATE, GraphST, and CCST was strongly dependent on the non-spatial clustering algorithm applied to the final embedding, since even the best-performing swap (maximum ARI) reduced the quality of the solution. On the other hand, applying dimension reduction like CCST during preprocessing helped to improve the performance of SCAN-IT, SEDR, and SpaceFlow which originally all used feature selection. At the same time, no individual component universally improved accuracy across methods (Supplementary Fig. 19), not even the neural network architecture that was critical to GraphST’s performance. Together, these results indicate that preprocessing and the final clustering stage account for a substantial share of performance variation, whereas architectural differences between neural networks contribute comparatively little.

**Figure 5:**
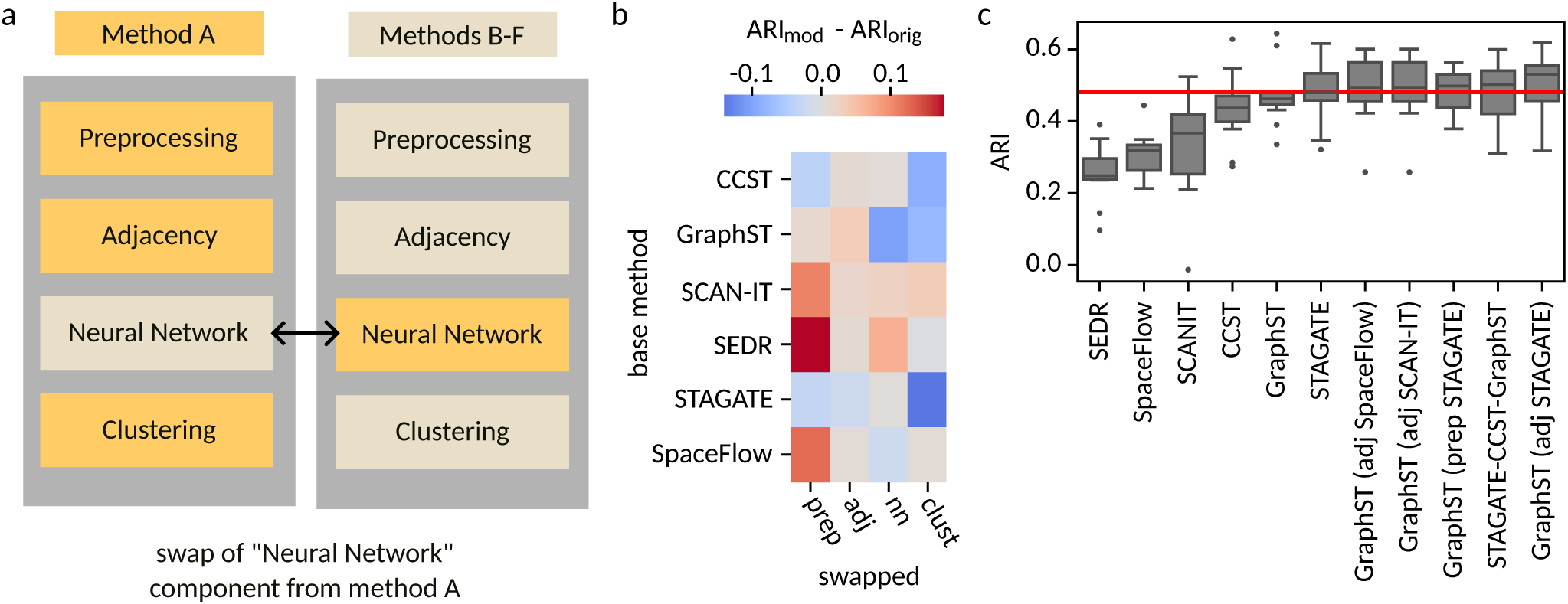
Ablation study of modularized neural network-based methods. **a**, Schematic of swapping one of four method modules. **b**, Maximum absolute ARI difference relative to original method after replacing, within each method, individual components for preprocessing (prep), adjacency calculation and graph construction (adj), neural network architecture and training (nn) and clustering of the learned embedding (clust). For each component swap, the mean ARI change across all 12 Visium-Maynard samples is reported. **c**, Performance of the base methods and the five best-performing recombined method configurations. The median performance of the best-performing base method (STAGATE) is indicated by a red line. For all best-performing combinations, clustering is performed using mclust. STAGATE-CCST-GraphST combines STAGATE for preprocessing, CCST for adjacency calculation, and GraphST for neural network training. Box plots: boxes show quartiles, whiskers extend to points within 1.5 interquartile lengths of upper and lower quartiles.

In addition to swapping individual components as above, we computed the clustering accuracy of module combinations when allowing all four components to vary simultaneously (Supplementary Fig. 20). Figure 5c shows the five combinations that achieved the highest median ARI. They all outperform the six original methods used in this experiment. Four of these combinations swap only a single component in GraphST, one combination mixes modules from three different methods. SpaceFlow and SCAN-IT use the 1-skeleton of the alpha-complex by default which slightly improves the (median) performance of GraphST when replacing its *k*-NN-based neighborhood calculation. Although the preprocessing in STAGATE is nearly identical to the one in GraphST, it slightly improves the median ARI achieved by GraphST. It omits the final normalization step after log-transforming the normalized counts. In summary, overall improvements from recombining modules were modest. The five highest-performing pipelines were all either small modifications of GraphST or incorporated the GraphST neural-network module.

### Runtime, memory usage, and usability

To assess runtime and memory usage, we used SRTsim [25] to generate simulated datasets with increasing numbers of cells. The number of cells ranged from 2,000 to 100,000, and for each dataset size we conducted three independent random trials (see Methods).

While most methods, with the exception of SpiceMix, complete within a few minutes on the smallest datasets, their scalability differs substantially (Supplementary Fig. 21). On the largest dataset comprising 100,000 cells, runtimes among the methods that completed successfully span several orders of magnitude, ranging from about one minute for the fastest method (BANKSY) to several hours and, in the case of MERINGUE, more than a full day. Memory usage shows a similarly wide spread. While all methods required less than 4 GB of memory on the smallest dataset, with several needing only a few hundred megabytes, memory consumption on the largest dataset increased substantially, ranging from a few gigabytes (e.g., TACCO and CellCharter) to approximately 200 GB (e.g., MERINGUE and GraphPCA). Several methods, including MNMST, DeepST, and ADEPT, exceeded the available memory limit already at smaller dataset sizes.

We grouped methods according to their growth rate in runtime and memory usage (Fig. 6a,b). Methods with slowly increasing runtimes can differ substantially in their absolute execution time, as illustrated by the red and orange groups. Notably, methods in the red group exhibit less favorable memory scaling than those in the orange group. Similarly, among methods with rapidly increasing runtimes, represented by the blue and green groups, methods with slightly longer absolute runtimes, shown in blue, scale worse in memory usage than somewhat faster methods shown in green.

**Figure 6:**
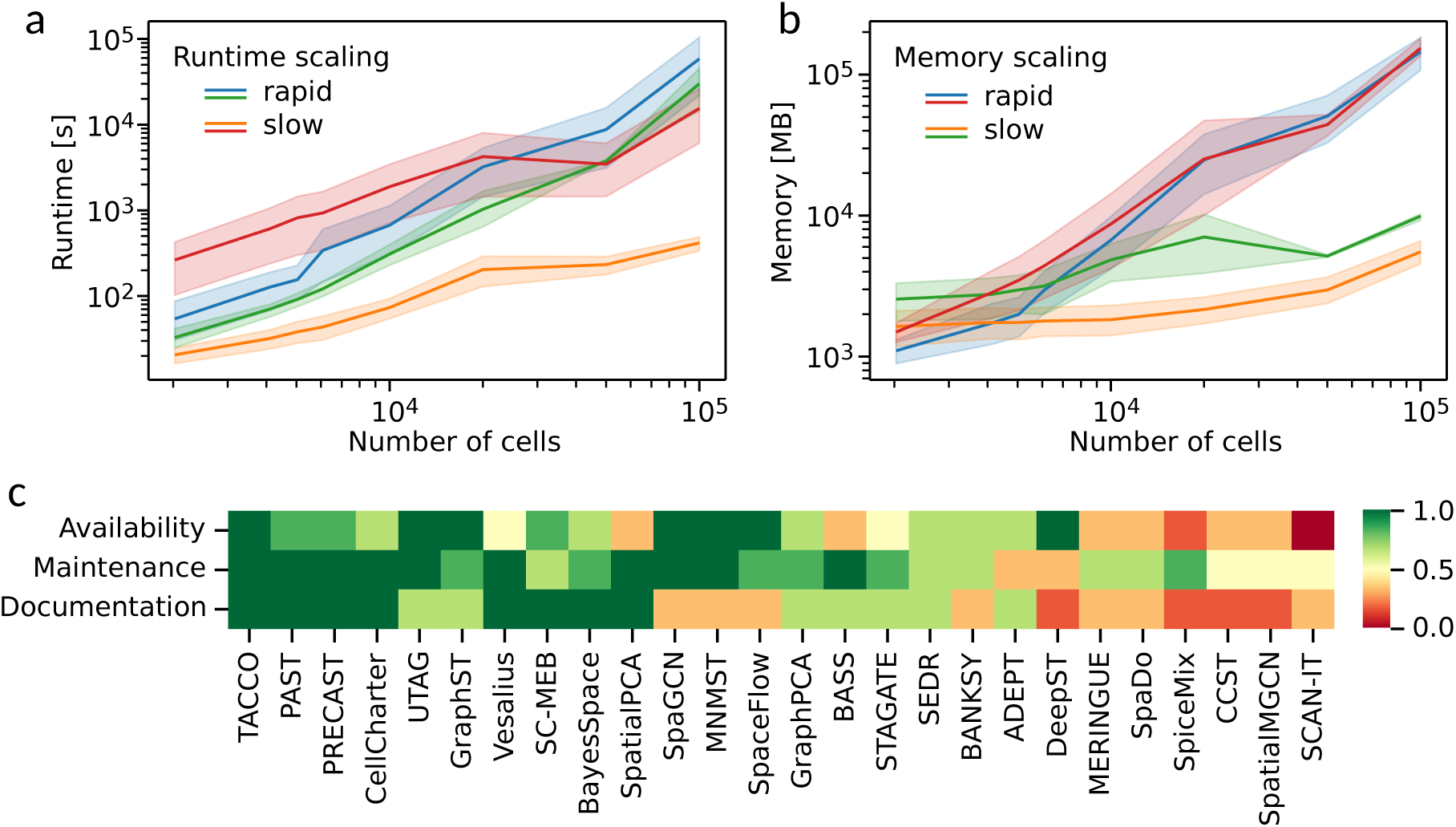
**a**, Runtime (seconds) and **b**, peak memory usage (MB) for all methods, grouped according to their scaling behavior. Solid line shows mean, spread shows 95% confidence interval. **c**, Usability scores based on availability (ease of installation), maintenance, and documentation (Methods). Scores range from 0 to 1, with higher values indicating better usability. Methods are ordered by their mean usability score.

Finally, we evaluated user-friendliness using a predefined checklist (Methods) and summarize the results in Fig. 6c. Only a few methods perform consistently well across all criteria, most notably TACCO, PAST, and PRECAST. While many tools satisfy several usability criteria, incomplete or fragmented documentation remains a common limitation across methods. In some cases, as also noted in Supplementary Note 3, discrepancies between the published descriptions and the released implementations further complicate method use and reproducibility, particularly for less experienced users.

## Discussion

Here, we present a comprehensive benchmarking study of spatial domain identification methods that moves beyond descriptive performance comparisons toward an explanatory understanding of method behavior. To achieve this, we implemented a modular and extensible pipeline for generating semi-synthetic spatial transcriptomics data, enabling systematic variation of both technological and tissue-inherent data characteristics. This framework allows us to connect method performance on real and semi-synthetic data, thereby identifying the factors that drive observed performance differences across technologies and datasets.

Previous benchmarking studies of spatial domain identification methods have largely focused on descriptive comparisons, reporting performance across individual datasets without attempting to explain underlying performance drivers [18–21]. In contrast, our benchmark explicitly adopts an explanatory paradigm, combining real and semi-synthetic data to identify the technological and biological factors that shape method behavior. This shift from “which method performs best” to “why methods perform differently” is essential for guiding both method development and method selection in a rapidly evolving technological landscape.

A central advantage of the semi-synthetic framework is access to a well-defined and controllable ground truth. While expert annotation of real datasets is an invaluable resource, a degree of uncertainty and arbitrariness in domain delineation is unavoidable. Moreover, despite the widespread use of the concept of spatial domains, there is currently no broadly accepted formal definition, which can lead to inconsistencies in both method development and expert annotation practices. Taken together, these considerations motivate the use of semi-synthetic data as a principled means to evaluate spatial domain detection methods across a broad and well-controlled range of domain and data characteristics.

As a starting point, we evaluated the performance of 26 methods and four baselines on seven publicly available real datasets, which revealed several technology-dependent patterns. Few methods strongly and consistently out-perform baseline performance on low-resolution data. In contrast, substantial improvements are observed on all high-resolution datasets, most notably those generated using MERFISH. We deliberately selected widely used and commercially available MERFISH and Visium datasets to represent opposite ends of the spectrum in both spatial resolution and the number of profiled genes. In direct comparisons, most methods exhibit a clear preference for one of these extremes, with this preference strongly correlating with the degree of spatial coherence observed in the inferred domain structures. These observations suggest that method performance is tightly coupled to underlying data characteristics, motivating a more targeted investigation of their individual effects.

To dissect these dependencies, we next turned to semi-synthetic data, which enabled systematic variation of both technology-inherent and tissue-inherent parameters. This analysis provides an insight into the drivers of method performance. We examined the impact of spatial resolution, gene panel size, and sparsity, as well as tissue properties such as transcriptional similarity between domains, cellular heterogeneity within domains, and domain size and shape. Across methods, cellular heterogeneity emerged as a dominant determinant of performance. Both purely transcriptionally informed baselines and many spatially informed tools showed substantial performance degradation as heterogeneity within domains increased. In contrast, methods that are resilient to transcriptional heterogeneity, including BASS, SpaceFlow, and SpaDo, consistently ranked among the top-performing approaches on high-resolution real datasets. Conversely, methods such as SpatialPCA, MNMST, and GraphST performed competitively on Visium data but were strongly affected by cellular heterogeneity and achieved only average performance on high-resolution datasets. These findings, summarized in Supplementary Table 1, both explain performance patterns observed in real data and reveal systematic dependencies that are not apparent from empirical datasets alone, underscoring the growing importance of spatially informed domain detection as spatial transcriptomics technologies approach cell-level and subcellular resolution.

Beyond comparisons of individual methods, our Snakemake-based benchmarking pipeline enables systematic analyses at the ensemble level. This allowed us to assess whether performance gains can be achieved by aggregating complementary method outputs rather than by further algorithmic specialization. Using a simple consensus strategy, we identified an approach that is highly competitive with, and in some cases outperforms, any individual method. Notably, the consensus analysis substantially improved performance on Visium datasets and remained competitive on high-resolution data. These results indicate that consensus approaches represent a promising strategy for improving both robustness and overall performance. In practice, their applicability may be limited by the runtime or memory requirements of individual methods included in the ensemble. However, restricting the consensus to smaller, carefully selected subsets of methods markedly reduces computational cost, making this strategy a viable complement to single-method analyses. Method subsets can be tailored to optimize overall performance or performance on specific technologies (Supplementary Fig. 22).

To further disentangle the sources of performance differences, we conducted an ablation study on a subset of neural network-based methods. This analysis probes the relative contributions of model architecture versus pipeline design. We found that core architectural innovations and carefully designed loss functions often have a limited impact on performance. In contrast, preprocessing strategies and final clustering choices exert a substantially stronger influence. Although several combinations of method components outperformed the best individual method in this ablation study, these improvements were modest, highlighting that improvements arise primarily from careful alignment and tuning of individual pipeline components rather than from architectural complexity in isolation.

We next examined stochastic variability of methods by developing a procedure that induces variation in random decisions that are otherwise implicit or fixed across methods, allowing us to quantify sensitivity to controlled stochastic perturbations. This analysis reveals that stochastic variability is widespread and largely independent of algorithmic class. Statistical models and neural network-based methods, as well as neural networks with different architectures, do not show consistent differences in stability. Instead, stochastic behavior is primarily shaped by preprocessing and postprocessing choices. For example, PCA-based dimensionality reduction tends to be more stable than variable gene selection, which is used by methods such as STAGATE, GraphPCA, and SCAN-IT. Methods that incorporate regularization in their loss functions, such as SpaceFlow, or apply stabilizing postprocessing steps such as PCA in CCST, also show reduced sensitivity to random initialization. Together, these results suggest that stochastic stability is driven more by pipeline design choices than by the underlying algorithmic framework.

As with any benchmarking effort, our study has limitations. We do not evaluate methods in a multi-slice domain identification setting, and all tools are assessed using default, technology-specific parameter settings as implemented or recommended by the authors (Methods). Performing comprehensive parameter sweeps is challenging because comparable hyperparameters are not defined consistently across implementations. For example, parameters controlling spatial smoothness or neighborhood size differ in meaning, scale, and default ranges between methods. In addition, as discussed in Supplementary Note 3, there are occasional inconsistencies between the mathematical descriptions in some manuscripts and their corresponding implementations, which further complicates standardized parameter tuning. We also exclude methods that require histological images, such as stLearn [52]. However, prior benchmarking studies suggest that histological information does not necessarily improve spatial domain detection performance [20]. More generally, all methods are provided only with expression and spatial location information, which may prevent tools capable of incorporating additional metadata (such as cell type annotations) from demonstrating their full potential. Finally, we apply all methods to all datasets irrespective of original design intent and rely on default parameterizations throughout, reflecting common usage patterns within the community.

From a user perspective, our results provide practical guidance for method selection. Several methods stand out for consistently strong performance across most evaluations, including BASS, SpaDo, PAST, GraphPCA, SCAN-IT, TACCO, SpaceFlow, and ADEPT. Among these, BASS emerges as a top recommendation in terms of pure performance. For users prioritizing ease of use and general usability, methods such as TACCO, CellCharter, PAST, and GraphST are attractive choices. For large datasets where runtime or memory constraints are limiting, TACCO, CellCharter, ADEPT, PAST, and SpaceFlow offer favorable scalability. Users should also take into account whether they require additional capabilities, such as support for multi-slice analysis, which is provided only by a subset of methods.

For method developers, several broader insights emerge. First, intercellular transcriptional heterogeneity within domains remains a major challenge for most existing tools, and this challenge is likely to become increasingly critical as spatial transcriptomics technologies continue to increase in resolution. Second, optimizing runtime and memory usage will be critical as dataset sizes grow. For developers of neural network-based methods in particular, our results emphasize the importance of carefully evaluating gains from architectural innovations independently of preprocessing and postprocessing choices. Finally, usability remains an underappreciated aspect of method development. Inadequate documentation and limited maintenance are common shortcomings, presenting clear opportunities for new methods to distinguish themselves.

In summary, this benchmark provides a comprehensive and explanatory evaluation of spatial domain identification methods, combining real and semi-synthetic data to uncover the mechanisms underlying method performance. By integrating performance, scalability, robustness, and usability, our study offers actionable guidance for method users and developers and provides a foundation for future methodological advances in spatial transcriptomics analysis.

## Methods

### Data acquisition and preprocessing

Table 1 shows all real datasets included in this study, along with selected metadata. Raw count matrices and spatial coordinates for all datasets were downloaded from the original data sources listed in Supplementary Note 2. We did not apply a unified preprocessing pipeline; instead, we followed the preprocessing procedures recommended for each individual method. All data were stored in a standardized csv format to ensure compatibility within the Snakemake pipeline. Manually curated domain annotations were obtained separately when they were not provided with the original dataset. Further details on the origin of the real datasets and their annotations are provided in Supplementary Note 2.

**Table 1:**
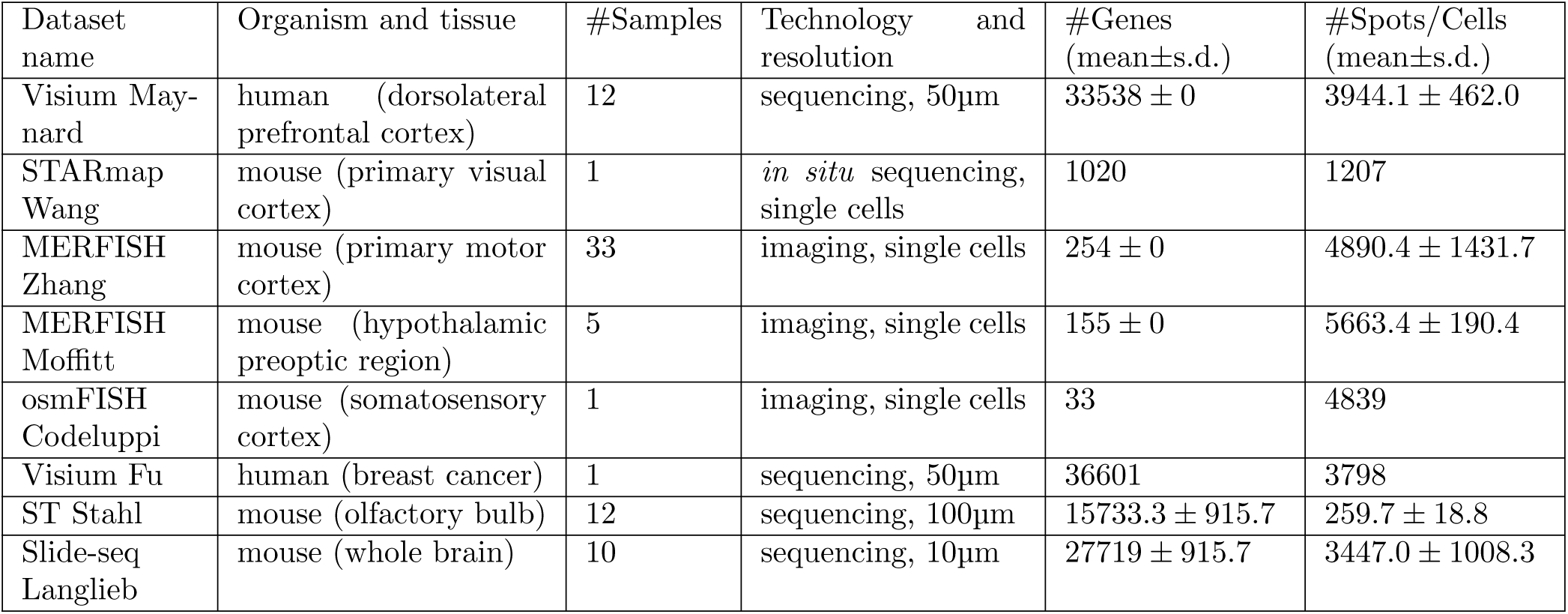
Metadata for all real datasets included in this study.

### Method parameter settings

We ran all methods using the default parameter settings recommended by the tool developers for the respective technology. Where defaults were explicitly specified in the code base, we applied them directly. Otherwise, we inferred recommended settings from tutorials or from the original publications. When technology-specific parameter recommendations were provided for only a subset of platforms, we followed those recommendations where applicable and defined analogous settings for technologies that were not explicitly covered. All scripts used to run the methods in this benchmark are included in the publicly available analysis framework (see Data and Code Availability).

### Methods included in the benchmark

Methods are described according to their broad categorization, shown in Fig. 1a.

#### Clustering-based methods

**BANKSY** computes a spatially weighted neighborhood gene-expression vector for each spot [41]. This vector is concatenated with the spot-level gene expression to form a neighbor-augmented feature vector in a joint product space. The relative contribution of neighborhood versus spot expression is controlled by a hyperparameter *λ*. Clustering is then performed in this augmented space using any conventional clustering algorithm.

**CellCharter** aggregates gene-expression features within neighborhoods of increasing size, defined using either Delaunay triangulation or *k*-NN graphs [46]. The resulting aggregated feature matrices are concatenated with the original gene-expression matrix, and the combined feature space is clustered using a Gaussian mixture model (GMM).

**MERINGUE** constructs a neighborhood graph using Voronoi tessellation, where two spots are considered adjacent if their Voronoi cells share a boundary [29]. A *k*-NN graph is then built in the space of the top principal components, and edge weights are assigned based on proximity in the neighborhood graph. Specifically, the weight between two nodes reflects the length of the shortest path between them in the graph. The resulting weighted graph is clustered using conventional graph-clustering methods.

**TACCO** constructs *k*-NN graphs from both the top expression principal components and the spatial coordinates, and then computes a weighted sum of the corresponding adjacency matrices [35]. The relative contributions of the two matrices are controlled by a hyperparameter. Leiden clustering is subsequently performed on the combined adjacency matrix. TACCO is primarily designed for annotation transfer and enables multi-slice analysis.

**UTAG** defines spatial adjacency by applying a distance threshold in physical space [28]. Expression values are then aggregated within each neighborhood by taking the mean across adjacent locations.

**SpaDo** is primarily designed for multi-slice spatial domain annotation [37]. It can either be provided with existing cell-type annotations or compute them directly from the data, using deconvolution for spot-level data. For each cell or spot, SpaDo computes the proportions of cell types within its neighborhood, defined using *k*-NN for cell-level data and a fixed spatial radius for spot-level data. A distance matrix between locations is then derived from these neighborhood cell-type proportion profiles using either the Jensen–Shannon divergence or the Manhattan distance, and hierarchical clustering is applied to identify spatial domains.

#### Statistical model-based methods

**BASS** uses a Bayesian hierarchical model that jointly infers both multi-slice cell-type and spatial-domain labels [33]. Gene expression is first projected onto the top principal components, which are modeled as multivariate normal with a covariance structure shared across cell types. Conditional on the spatial-domain labels, cell types follow a categorical distribution. A Potts prior is imposed on the spatial-domain labels to encourage spatial smoothness. In this framework, spatial domains are therefore modeled as mixtures of cell types, rather than being inferred directly from gene expression.

**BayesSpace** is a fully Bayesian method that incorporates spatial information through a hidden Markov random field (HMRF) prior [45]. The top principal components of gene expression are modeled as a multivariate Gaussian distribution with covariance shared across domains. A Potts prior is imposed on the domain labels, penalizing assignments in which neighboring spots belong to different domains. The strength of this spatial smoothing is governed by a hyperparameter *γ*.

**PRECAST** combines a hidden Markov random field (HMRF) model with a Potts prior and an intrinsic conditional autoregressive (CAR) model to jointly embed and cluster spatial transcriptomics data [50]. It is particularly designed to align embeddings across multiple tissue slices.

**SC-MEB** implements a Bayesian HMRF model, but replaces the fully Bayesian inference of BayesSpace with an empirical Bayes approach based on maximizing a pseudo-likelihood [49]. A key contribution of SC-MEB is the use of a modified Bayesian information criterion (MBIC) to automatically determine the optimal number of clusters. In addition, the spatial smoothing parameter *β* is selected via grid-search optimization.

**SpiceMix** integrates a spatial prior implemented through an HMRF with a non-negative matrix factorization (NMF) model of gene expression [27]. Gene expression is decomposed into a mixture of metagenes, whose spatial affinities are inferred via the spatial prior. The learned metagenes are then clustered using conventional clustering methods to define spatial domains.

**GraphPCA** incorporates spatial information by adding a graph-based regularization term to the standard PCA objective [39]. This yields a closed-form solution that produces latent factors corresponding to spatially informed principal components. The resulting embeddings are then clustered using conventional algorithms.

**SpatialPCA** extends probabilistic PCA by incorporating spatial information through a Gaussian kernel–based covariance structure [32]. Model parameters are estimated via maximum likelihood, and the resulting latent factors are clustered using conventional algorithms.

#### Neural network-based methods

The neural network–based methods are described individually below and placed within a unified conceptual frame-work in Supplementary Note.

**MNMST** defines spatial adjacency using pointwise mutual information [31]. Gene expression is first augmented with local-neighborhood features using BANKSY. An adjacency matrix is then learned from the augmented expression profiles via sparse self-representation learning. MNMST jointly estimates shared cell features and an affinity graph, and the resulting embeddings are clustered using conventional clustering algorithms.

**CCST** constructs a spatial adjacency graph based on a user-defined radius around each spot or cell, with a hyperparameter controlling the relative weight of spatial information in the embedding [30]. It applies Deep Graph Infomax (DGI) to this weighted graph, generating corrupted graph samples by permuting edges [53]. (DGI is described in more detail in Supplementary Note.) The learned DGI embeddings are then reduced in dimensionality using PCA and clustered using conventional methods.

**SCAN-IT** defines spatial adjacency using the *α*-complex, which is derived from the Voronoi tessellation and a distance threshold [40]. A graph convolutional network (GCN) is then trained using Deep Graph Infomax (DGI), where corrupted samples are generated by permuting node features. A consensus distance matrix is constructed from independently trained DGI models, and the final low-dimensional representation is obtained using metric multidimensional scaling (MDS). This embedding is subsequently clustered using conventional clustering algorithms.

**SpaceFlow** constructs a spatial adjacency graph using an *α*-complex approach [34]. It applies the Deep Graph Infomax (DGI) framework, using node permutation to generate corrupted graphs. In addition, SpaceFlow introduces a user-specified spatial regularization term in the loss function to encourage spatial consistency. The learned embeddings are then clustered using conventional clustering algorithms.

**GraphST** constructs a spatial *k*-NN graph and generates a corrupted counterpart by randomly permuting node features [44]. It employs a GCN-based encoder–decoder architecture with an objective function that combines a self-reconstruction loss with a DGI-inspired contrastive loss applied to both the original and corrupted graphs. The reconstructed gene-expression profiles are then clustered to identify spatial domains.

**STAGATE** constructs a radius-based binary spatial adjacency matrix [38]. For grid-based, low-resolution data, the graph can optionally be pruned based on a coarse gene-expression preclustering obtained using the Louvain algorithm. A graph-attention autoencoder is then used to learn spatially informed embeddings by minimizing a reconstruction loss. The resulting embeddings are clustered using conventional clustering algorithms.

**ADEPT** first trains a graph-attention autoencoder on the spatial *k*-NN graph to learn node embeddings, which are clustered (via GMM) to define preliminary domains [51]. Differentially expressed genes (DEGs) are then selected based on non-zero-rate thresholds, and multiple DEG-based expression matrices are imputed to remove dropouts. These imputed DEG matrices are averaged and fed back into the same graph-attention autoencoder, whose final embedding is clustered again to define the spatial domains.

**PAST** uses a Bayesian neural network (BNN) to optionally incorporate reference gene-expression data alongside standard fully connected networks (FCNs) [47]. Its objective function combines a reconstruction loss, a Kull-back–Leibler divergence term to a standard normal prior, the BNN loss, and a metric-learning loss. The final embeddings are obtained through self-attention and FCN layers and are subsequently clustered to define spatial domains. To enable scalable training on large datasets, PAST employs a ripple-walk sampling strategy for mini-batching.

**DeepST** constructs a modified gene-expression matrix that incorporates information from spatial neighbors, weighted by their expression correlation and, optionally, morphological similarity [36]. A spatial *k*-NN graph is also built from the coordinates. DeepST then applies both a denoising autoencoder and a variational graph autoencoder (VGAE), optimizing a reconstruction loss together with a Kullback–Leibler divergence term. The resulting embeddings are clustered using conventional algorithms to define spatial domains.

**SEDR** applies a masked self-supervised scheme in which the expression profiles of a random subset of spots are replaced by a learnable mask vector, and learns a low-dimensional expression embedding with a deep autoencoder from this masked matrix [48]. A variational graph autoencoder (VGAE) with a GCN encoder takes the spatial *k*-NN adjacency and the expression embedding as node features to obtain a spatial embedding. The two embeddings are concatenated into a joint latent representation that is used to reconstruct both gene expression and the spatial adjacency, by minimizing the cross-entropy loss between the learned and true adjacency matrices together with a KL divergence term to a Gaussian prior. The final latent representation is clustered using mclust to define spatial domains.

**SpatialMGCN** constructs a binary spatial adjacency graph using a radius-based criterion and, in parallel, builds a *k*-NN feature graph from the gene-expression matrix using cosine distance [42]. A multi-view GCN encoder then performs convolutions on the spatial and feature graphs both separately and jointly, and an attention mechanism is applied to integrate the resulting embeddings. The decoder models gene expression using a zero-inflated negative binomial (ZINB) distribution, and the reconstruction loss is given by the negative log-likelihood of the ZINB model. In addition, SpatialMGCN includes a spatial-regularization loss that penalizes large embedding distances between spatially neighboring spots or cells

**SpaGCN** constructs a complete, weighted, undirected graph of spots or cells, where edge weights are computed from the Euclidean distance in physical space using a Gaussian kernel; optionally, morphological features can be incorporated as an additional spatial dimension [54]. A single graph-convolutional layer is then applied to learn embeddings that integrate both the expression matrix and the spatial graph structure. Cluster centroids are initialized from this embedding using Louvain clustering, and the cluster assignments are subsequently refined through an iterative optimization strategy.

**Vesalius** embeds the gene-expression matrix into a three-dimensional latent space using UMAP and interprets the three coordinates as RGB channels to construct an image representation of the tissue [55]. This image is then iteratively smoothed and segmented. For each resulting segment, spots within a specified capture radius are pooled to define spatial domains.

#### Baseline clustering methods

We included two baseline methods that do not exploit spatial information and instead cluster cells or spots purely based on gene expression. For this purpose, we used two widely adopted clustering implementations for single-cell data: Leiden clustering as implemented in scanpy [22] and Seurat [23].

In addition, we constructed naïvely spatially aware baselines by post-processing the scanpy– and Seurat-derived clusters using a spatial smoothing procedure adapted from SpaGCN [54]. In this procedure, spot labels are reassigned by majority vote over the local spatial neighborhood defined by the *n* nearest neighbors, with *n* = 4 for ST-Stahl, *n* = 6 for Visium datasets, and *n* = 5 for other technologies. These values account for the differing spatial layouts of the respective platforms.

### Consensus approach

We implemented a simple consensus strategy that aggregates the label outputs of multiple methods. For each spot, the consensus label is obtained by taking the mode of the predicted labels across methods. To make this approach meaningful, the label assignments from different methods must first be mapped to a common label set. Since ground-truth labels are available for all samples in this study, we aligned the predicted cluster labels to the ground truth by solving a maximum-weight bipartite matching problem, implemented via the linear-sum assignment solver in the scipy package [56].

### Accuracy and spatial coherence metrics

We used the Adjusted Rand Index (ARI) as the primary accuracy-based evaluation metric [57–59]. The ARI is the chance-corrected version of the Rand Index (RI), defined as

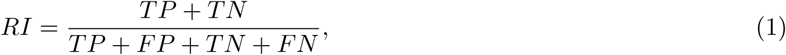

where *TP* denotes the number of true positives, *FP* the number of false positives, *TN* the number of true negatives, and *FN* the number of false negatives. The ARI corrects for the non-zero expected value of the RI under random label assignments and ranges from 1 (perfect agreement) to –1 (maximal disagreement), with 0 indicating performance equivalent to random assignment.

To quantify the spatial coherence of clustering results, we used the Percentage of Abnormal Spots (PAS), adapted from an image analysis metric [32]. A spot (or cell) is considered abnormal if its cluster label differs from the label of more than half of its *k* = 10 nearest spatial neighbors. Lower PAS values indicate higher spatial coherence and reduced visual noise.

We further assessed domain-specific accuracy using the F-score, the harmonic mean of precision and recall, defined as

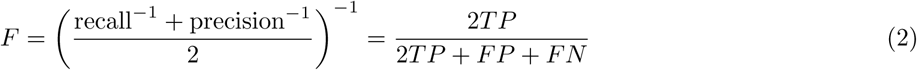

In practice, the F-score is computed in *scikit-learn* as a collection of binary classification problems evaluated over all pairs of domains [60]. When two domains receive identical labels and are therefore indistinguishable, the overall F-score may still remain relatively high because large correctly separated domains can dominate the score. To explicitly quantify how well individual domain pairs can be distinguished, we defined a pairwise domain indistinguishability metric, which we term confusion. Let *N^j^* denote the number of spots belonging to ground truth domain *i* that are assigned method-predicted label *j*. We define

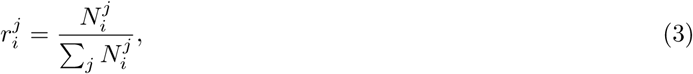

that is, 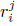 is the proportion of spots from ground truth domain *i* assigned to label *j*. The confusion between domains *a* and *b* is then given by

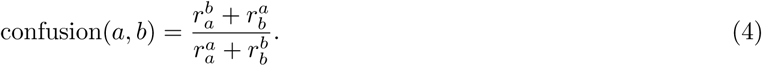

A value of 0 indicates that domains *a* and *b* are well separated, whereas a value of close to 1 indicates strong mixing or indistinguishability. To obtain a domain-level confusion score, we evaluate the maximum of pairwise confusion values involving that domain.

### Evaluation of stochastic variability

We evaluated the stochastic variability of each method by rerunning it multiple times on the same data. Several methods internally fix the random seed controlling stochastic components, which results in identical outputs across repeated runs. To uncover variability even in such cases, we perturbed the input order of the data in each repeated execution.

For each sample, multiple perturbed instances were generated by randomly permuting the spot (or cell) indices. The permuted indices were then used to reorder the count matrix, spatial coordinates, and labels in a consistent manner. This procedure leaves the data values unchanged and alters only the ordering of inputs, which may expose latent stochasticity.

Stochastic variability was assessed across multiple samples from two datasets, namely Visium-Maynard and MERFISH-Zhang. To account for inter-sample differences in baseline performance, ARI values were linearly rescaled on a per-sample basis such that the sample-specific median matched the dataset-wide median, while preserving the relative variability within each sample.

### Generation of semi-synthetic data

We generated semi-synthetic spatial transcriptomics dataset using the following procedure. The simulation code is available as described in the Data and Code Availability section.

#### Generation of spatial geometries and ground-truth domains

To avoid shape-specific biases from influencing method comparisons, we generated semi-synthetic tissues with multiple domain geometries. The selected layouts represent archetypal tissue organizations frequently observed in spatial transcriptomics data: laminar structures, single circular domains, concentric circular domains of varying size, and a complex interlocking configuration (visualized in Fig. 4c). For analyses focusing on technological parameters (e.g. resolution, panel size, sparsity) or count-level perturbations independent of geometry, performance metrics were averaged across all layouts to mitigate potential shape-driven bias.

For all tissue configurations except layered arrangements, initial domain prototype points are generated using synthetic data generators from the scikit-learn datasets module [60]. Spatial spot locations are then randomly sampled from a uniform distribution using NumPy [61]. Unless stated otherwise, we used *n* = 4,000 spatial locations throughout this study. Domain labels are assigned to spots by transferring labels from the prototype points in Euclidean space. Spots which are not the nearest neighbor of any prototype are assigned to a background class, and isolated outlier spots are reassigned to the majority label within their local neighborhood to promote spatial coherence. For layered configurations, domain labels are defined directly on the sampled spot locations by specifying linear boundaries that partition the tissue into distinct layers.

#### Cell type selection and count assignments

Single-nucleus mouse brain data were downloaded from docs.braincelldata.org/downloads, together with fine-grained cell-type annotations [24]. We selected five cell types (Table 2) whose pairwise distances did not exceed 0.01 units according to [24] (Supplementary Fig. 12) and that each contained more than 1,500 cells. Cell types from the single-nucleus data were then mapped to domain labels, and count vectors of individual nuclei from these cell types were assigned at random to previously generated spatial locations within the corresponding domains. By varying the random seed used for count assignment, we generate multiple replicates of each synthetic tissue. Specifically, five replicates are created for all samples used to evaluate the effects of technological parameters (resolution, number of genes, and sparsity), count-level perturbations, and domain size.

**Table 2:**
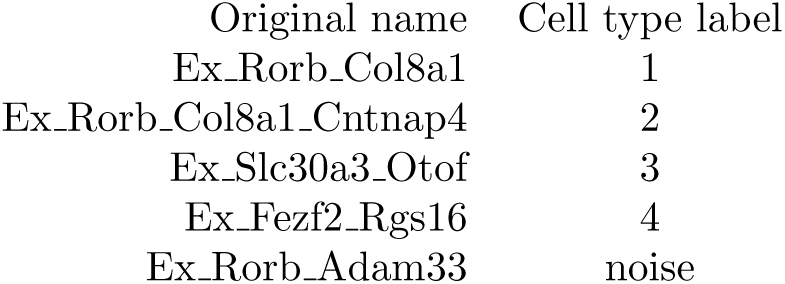
Cell types selected from [24] and their corresponding labels used in this study. Ex_, excitatory neuron.

| Original name | Cell type label |
| --- | --- |
| Ex_Rorb_Col8a1 | 1 |
| Ex_Rorb_Col8a1_Cntnap4 | 2 |
| Ex_Slc30a3_Otof | 3 |
| Ex_Fezf2_Rgs16 | 4 |
| Ex_Rorb_Adam33 | noise |

#### Implementation of perturbations

Technological perturbations were applied to the generated data to mimic variation in gene coverage, spatial resolution, and sparsity. These were implemented as follows:

- **Downsampling of genes:** The target number of genes was defined as a proportion of the total number of genes in the single-nucleus dataset (approximately 21,000). The specified number of genes was selected uniformly at random and retained, while all remaining genes were removed.
- **Aggregation into larger spots:** To simulate lower spatial resolution, continuous spot coordinates within a tissue of size 100*×*100 (arbitrary units) were binned to coarser grids with side lengths of 0.5, 1, 2, 3, 5, 7, and 10. Spots mapping to the same aggregated location were merged: domain labels were assigned by majority vote (mode), and expression counts were averaged across all spots within each aggregated location.
- **Increase of sparsity:** The initial sparsity of the count matrix was approximately 85%. We increased sparsity up to 99% by randomly setting additional non-zero entries to zero. For each target sparsity level, the required number of entries to zero-out was computed based on the desired zero proportion, and the corresponding gene–cell pairs were sampled uniformly at random.

Biological perturbations of type I and type II were generated by modifying the composition of domain-specific expression profiles:

- **Type I perturbation:** A proportion of noise expression was added to the domain-specific expression profiles (i.e., gene expression from the noise cell type, see Table 2). The mixing proportion was varied from 0 to 100%. For each gene count, the observed value was replaced by a convex combination of the domain-specific and noise expression values.
- **Type II perturbation:** A subset of nuclei in each domain was replaced by nuclei drawn from the noise cell type (see Table 2). As in the type I perturbation, the replacement proportion was varied from 0 to 100%.

#### Assessment of shape-specific effects

For the analysis of shape-specific effects (Fig. 4c), we generated samples under all possible permutations of cell-type-to-domain assignments, ensuring that observed performance differences are not driven by a particular alignment of transcriptional similarity and spatial configuration.

Because the original layouts differ in the number of domains, direct comparison would introduce a structural bias. To ensure comparability across shapes, layouts containing a larger number of domains were redefined to match a common domain count. This normalization step ensures that differences in performance reflect geometric properties of the layouts rather than trivial differences in clustering complexity induced by varying domain numbers.

### Modularization of neural-network-based methods

We decomposed the source code of 6 neural-network-based methods, STAGATE, SCAN-IT, SEDR, CCST, GraphST and SpaceFlow, into three independent modules: preprocessing, neighborhood graph construction, and neural network architecture and training (Supplementary Table 2). For some methods, these modules were already implemented as separate functions and could be reused directly. In others, multiple steps were tightly coupled within single functions; in these cases, we disentangled the steps and extracted them into separate functions to enable modular reuse. In addition, whenever parameters were hard-coded to constant values within the original implementations, we refactored the code to expose these quantities as explicit function arguments and initialized them using the values recommended by the respective authors. Apart from the shared AnnData [62] input format, a Python package for annotated data matrices, there was no consistent standard for internal data structures (e.g. edge lists vs. adjacency matrices, sparse vs. dense matrices). We therefore standardized the representations using a sparse adjacency matrix format throughout and, where necessary, transformed the inputs and outputs of each module to ensure compatibility within our framework. For each resulting pipeline combination, we clustered the learned embeddings using k-means and *mclust*, as implemented in *scikit-learn* [63] and *R-mclust* [64], respectively. The refactored modular framework is publicly available as described in the Data and Code Availability section.

### Measuring runtime and memory usage

To systematically evaluate the computational requirements and practical usability of spatial domain detection methods, we generated synthetic datasets of varying sizes using the SRTsim package [25]. We simulated datasets containing 2,000, 4,000, 5,000, 6,000, 10,000, 20,000, 50,000, and 100,000 cells, with three replicates per size (random seeds 1, 2, and 3). Cells were randomly distributed within a square tissue layout and partitioned into four stripe-like spatial domains defined by x-coordinate thresholds at 0.25, 0.50, and 0.75. Domains differed in expression with fold changes of 1, 3, 2, and 4, respectively. Gene expression consisted of 250 high-signal, 250 low-signal, and 500 background genes (1,000 genes total), with controlled sparsity (5% zero inflation) and overdispersion (dispersion parameter 0.5).

All methods were executed within a Snakemake workflow, which automatically recorded the wall-clock time and maximum resident set size for each run. Experiments were performed on an AMD Ryzen Threadripper 3990X 64-core processor (4.3 GHz) equipped with 256 GB RAM.

### Usability evaluation

We evaluated the usability of each method based on three main criteria: availability (including ease of installation), maintenance (including code quality), and documentation (including tutorials). Each criterion was further divided into three minimally subjective sub-criteria that capture different aspects of the user experience, loosely following the framework of Rao *et al.* [65]. For each sub-criterion, methods were manually assigned a score of 0 (not fulfilled), 0.5 (partially fulfilled), or 1 (fully fulfilled). If a sub-criterion could not be assessed for a specific method, its score was omitted from the evaluation. Scores were averaged within each criterion, and the full list of criteria and method-level evaluations is provided in Supplementary Table 3.

## Data and code availability

Details on the availability and download sources of all real datasets used in this study are provided in Supplementary Note 2. The full analysis framework, including the extensible Snakemake workflow used for benchmarking and the modularized neural-network components described in the Methods, is publicly available at https://github.com/canzarlab/spatial-domain-analysis-framework. The repository includes scripts required to reproduce the analyses presented in this study, together with documentation describing how additional methods, modules, or datasets can be integrated into the framework. In particular, it also provides the refactored preprocessing, neighborhood graph construction, and neural network training modules that enable plug-and-play recombination across methods. The code used to generate the domain-aware semi-synthetic datasets is available at https://github.com/canzarlab/spatial-domain-simulator, including a full tutorial.

## Acknowledgements

The work was supported by German Research Foundation (DFG) grant CRC TRR338, project number 452881907, project Z03 (J.K. and S.C.) and an Else-Kröner-Fresenius-Stiftung starting grant 2019 A70 (J.K.). D.M. was sup-ported by the National Recovery and Resilience Plan 2021–2026 (NPOO) under the project ‘Advanced Algorithms and Optimization Models Supported by Mathematical Theory – OptimaAI’ (581-UNIOS-54). V.H.D. gratefully acknowledges the hospitality and financial support provided by Vietnam Institute for Advanced Studies in Mathematics (VIASM) through the *IAPB* program. A.D. was supported by the Graduate School of Quantitative and Molecular Biosciences Munich.

