## Supplementary Material for "An explanatory benchmark of spatial domain detection reveals key drivers of method performance"

#### Supplementary Figures

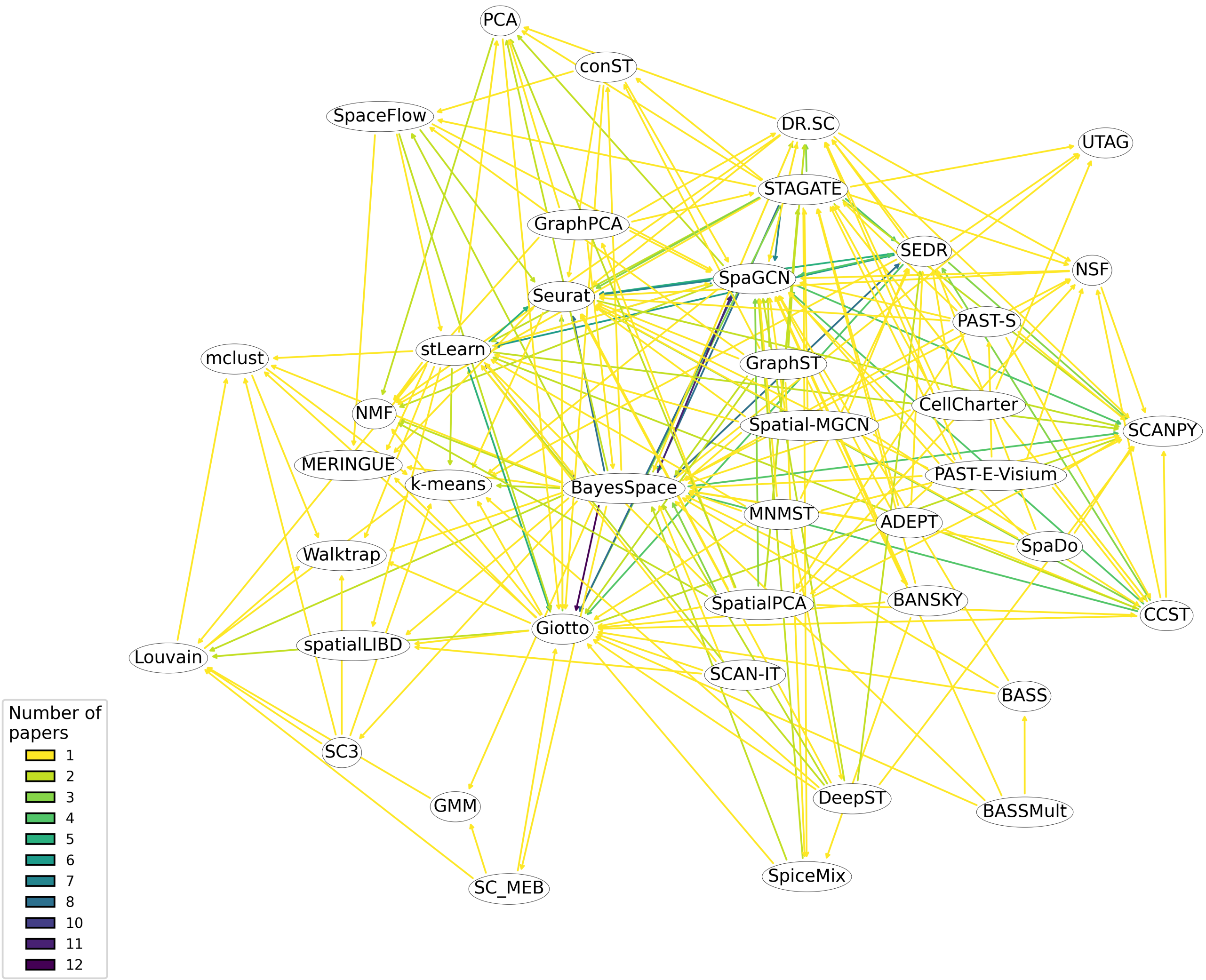


Supplementary Figure 1: Evaluation of reported method performance comparisons on the Visium Maynard dataset, manually extracted from the original publications of all methods included in this benchmark. Pairwise performance claims are represented as edges between methods, weighted by the number of publications reporting a given comparison. Edges are directed from the better-performing to the worse-performing method.

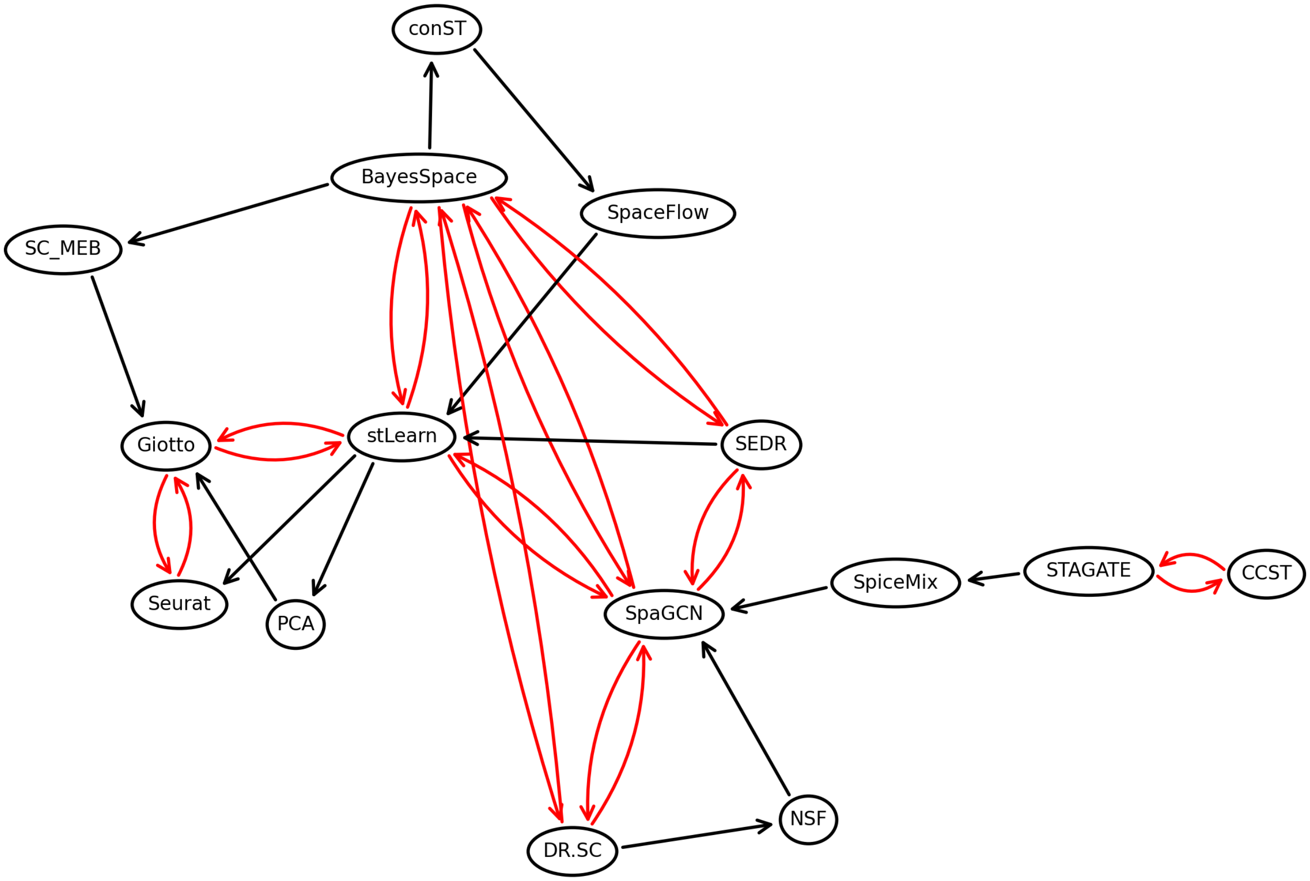


Supplementary Figure 2: Transitive reduction of the graph of reported method performance comparisons shown in Supplementary Fig. 1. Direct contradictions between reported performance claims are highlighted in red.

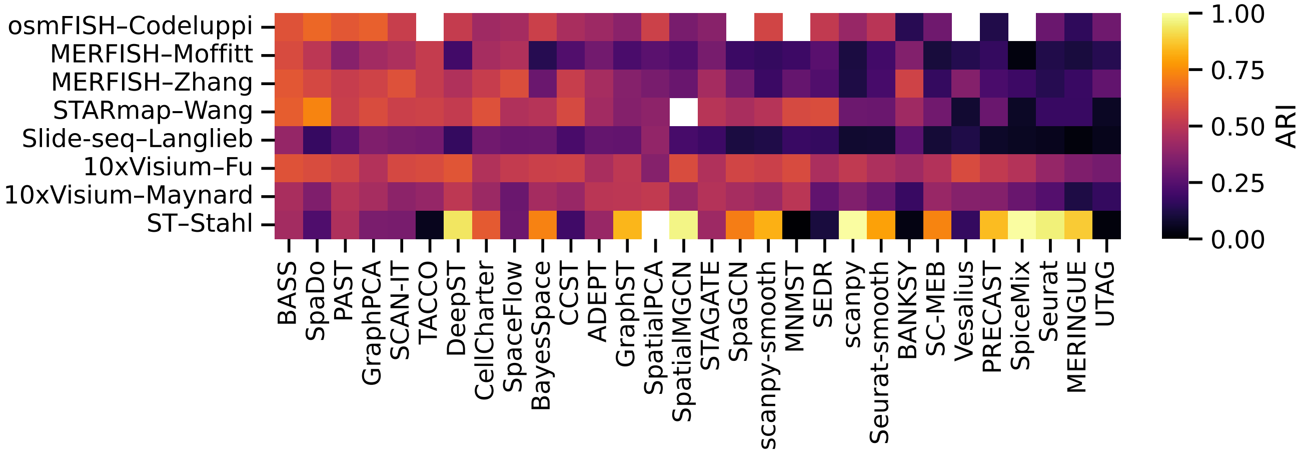


Supplementary Figure 3: Adjusted Rand indices (ARI) of all methods across all real datasets, aggregated as the mean over samples for multi-sample datasets. Methods are ordered by their mean standardized ARI. Blank squares indicate missing values, corresponding to cases where no clustering output was generated.

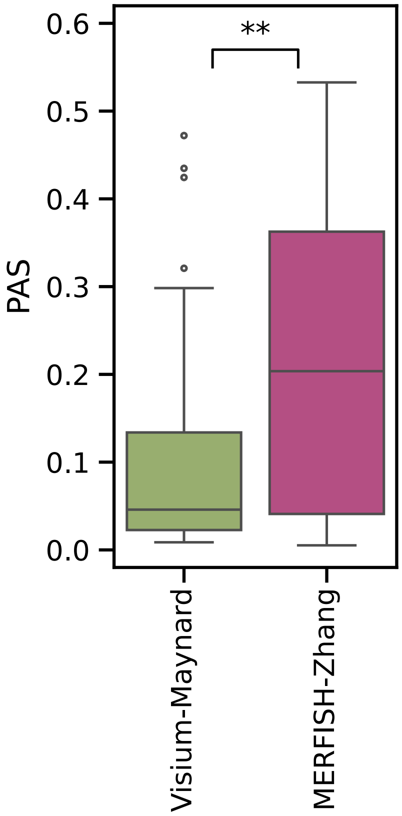


Supplementary Figure 4: PAS values for the inferred domains of all methods across all samples from the Visium Maynard and MERFISH Zhang datasets. Significance was assessed using a one-sided t-test ($p < 0.01$).

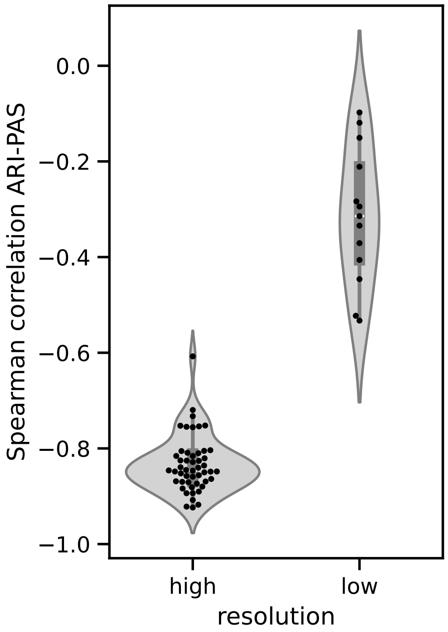


Supplementary Figure 5: Spearman correlation between ARI and PAS across all methods, stratified by data resolution. Visium datasets are classified as low resolution, and all other technologies as high resolution. The ST-Stahl dataset was excluded from this analysis.

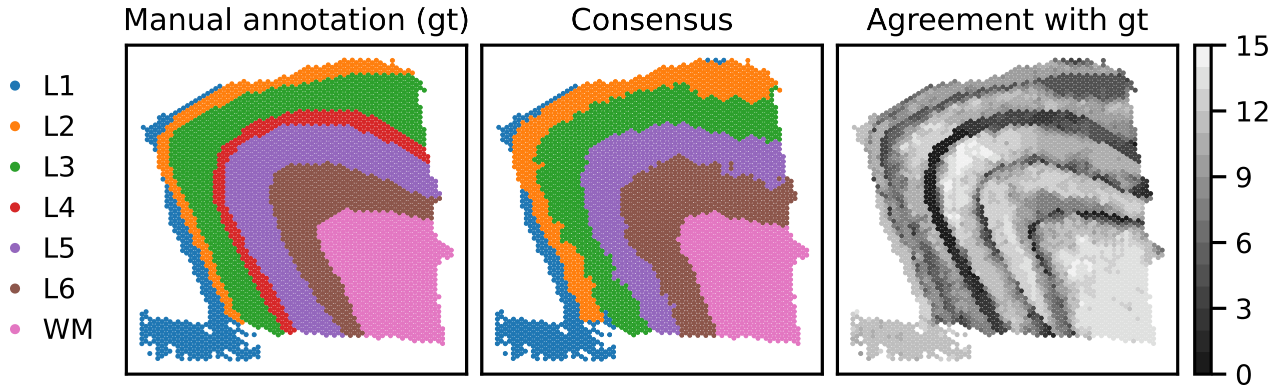


Supplementary Figure 6: Comparison of manually annotated and inferred consensus domains on slice 151675 of the Visium-Maynard dataset. Left, Manual gold standard annotation taken from the original publication [1]. Middle, Consensus of domains inferred by 15 top-performing methods (Methods). Right, Number of methods that agree on each spot label with the manual annotation.

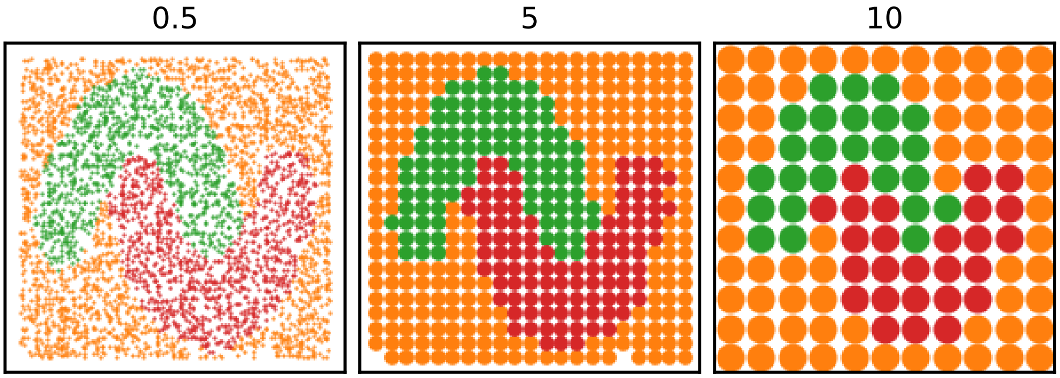


Supplementary Figure 7: Examples of semi-synthetic tissue maps at different spatial resolutions. Samples are generated by binning original cell locations onto grids, here with side lengths 0.5, 5, and 10.

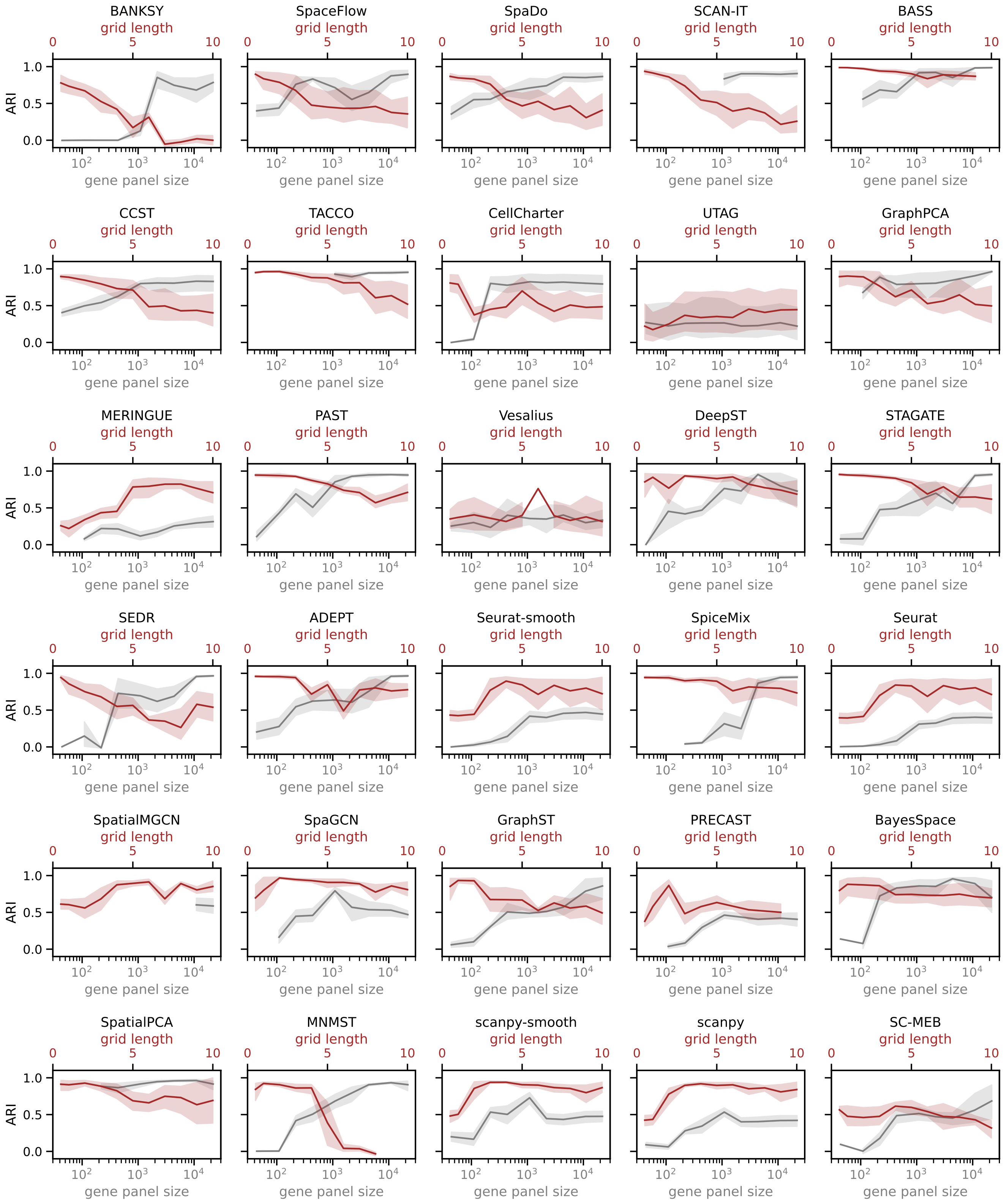


Supplementary Figure 8: Performance as a function of resolution (grid side length) and gene panel size, measured by ARI. Methods are ordered according to their relative performance on real MERFISH and Visium data (see Fig. 2b), with methods performing better on MERFISH shown toward the top left and those performing better on Visium toward the bottom right. Within each plot, moving from left to right along the $x$-axes corresponds to a transition from a MERFISH-like setting (small gene panel, no grid aggregation) to a Visium-like setting (full-transcriptome profiling, large grid side length). Variance reflects differences in tissue layouts (domain shapes) in the semi-synthetic data. Gray lines terminate early for methods that did not produce outputs at smaller gene panel sizes.

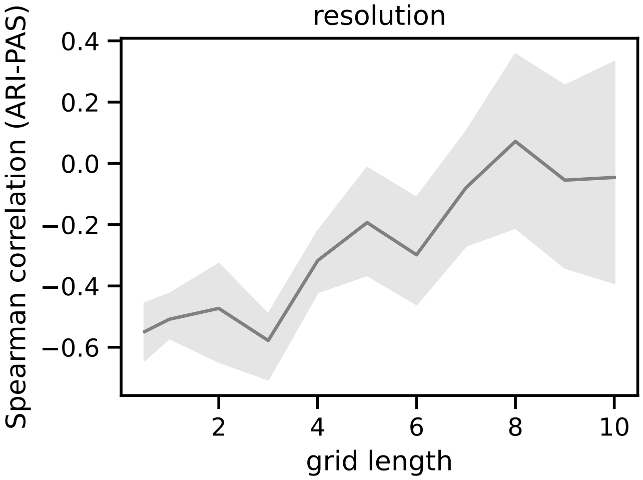


Supplementary Figure 9: Spearman correlation between ARI and PAS as a function of grid side length, which controls spatial resolution in the semi-synthetic data.

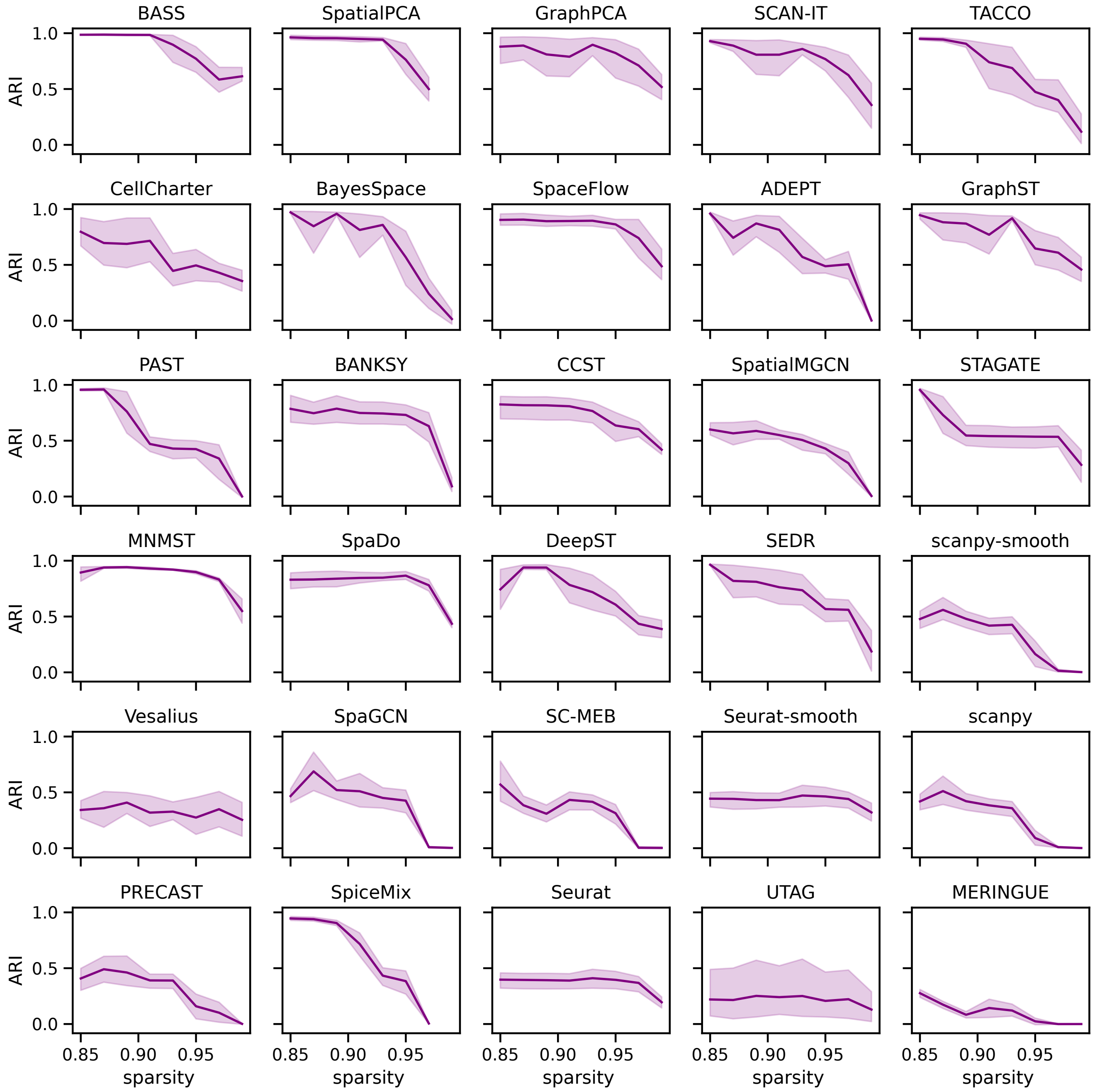


Supplementary Figure 10: Performance as a function of data sparsity, measured by ARI. Methods are ordered as in Supplementary Fig. 8. Variance reflects differences in tissue layout (domain shapes).

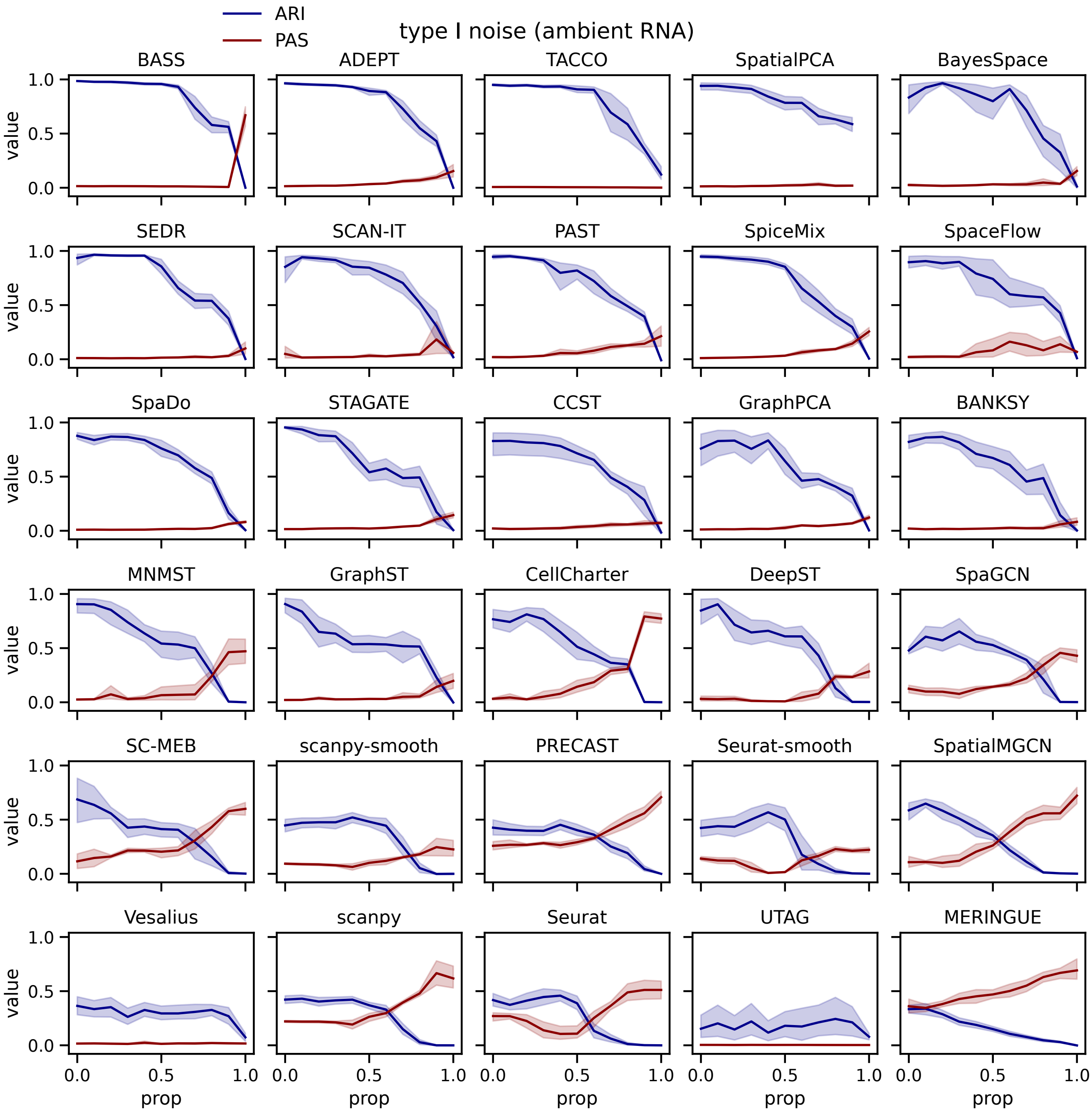


Supplementary Figure 11: Dependence of ARI and PAS on the level of type I perturbation (ambient RNA). The noise proportion (prop) denotes the percentage of each count contributed by a noise cell type (see Methods). Variance reflects differences in tissue layout (domain shapes) and five random seeds used for cell assignment. Methods are ordered by the area under the ARI curve.

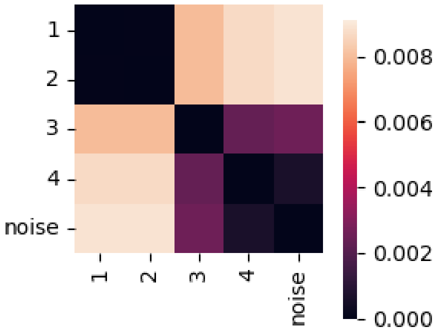


Supplementary Figure 12: Pairwise distances between the cell types selected for semi-synthetic data generation. Distances are reported in the same units as in the dendrogram of Langlieb *et al.* [2]. Cell types 1-4 denote the primary types used for data generation, with the noise cell type additionally included in some variations (see Methods).

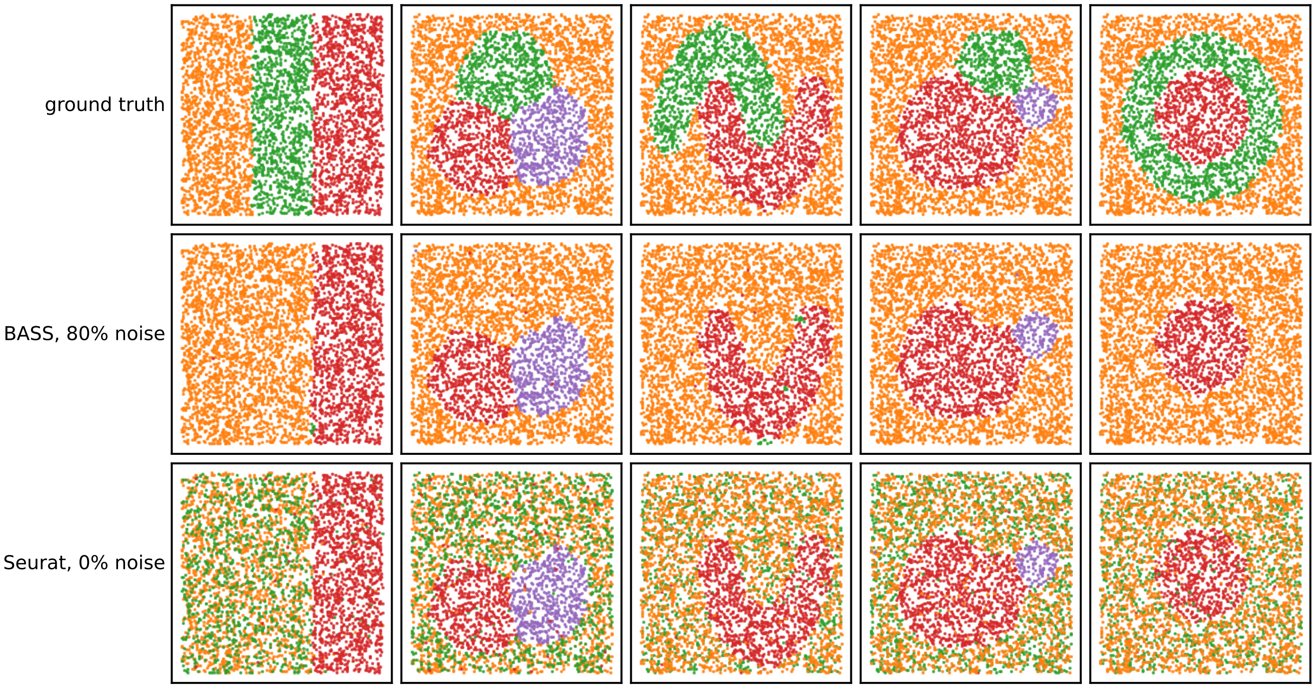


Supplementary Figure 13: Example tissue maps for different domain shapes (columns). The top row shows the ground truth domains, the middle row shows domains inferred by BASS after 80% type I perturbation, and the bottom row shows domains inferred by Seurat without perturbation. The green and orange domains contain cell types 1 and 2, respectively, which are the two most similar cell types (see Supplementary Fig. 12).

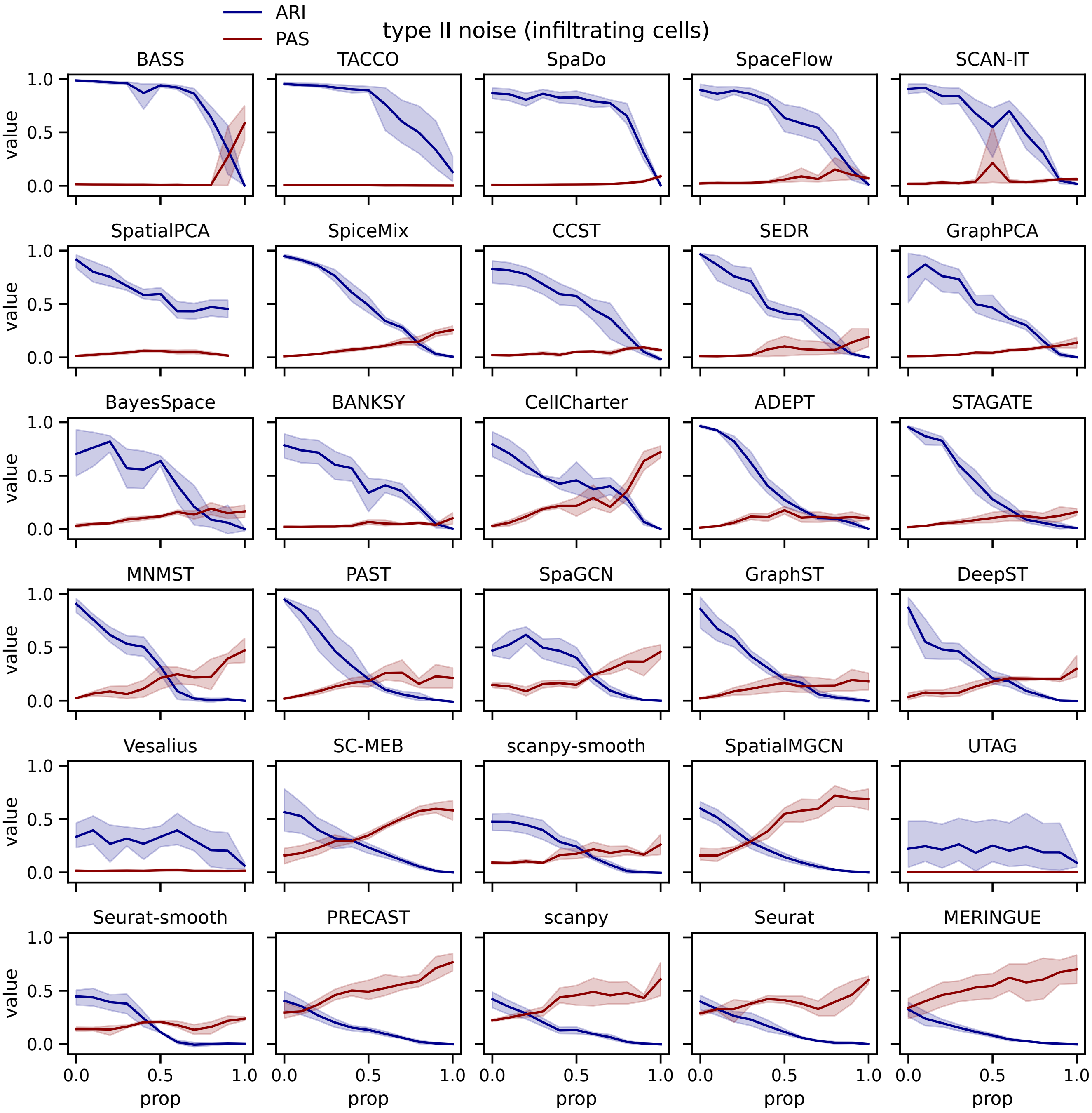


Supplementary Figure 14: Dependence of ARI and PAS on the level of type II perturbation (infiltrating cells). The noise proportion (prop) denotes the percentage of infiltrating noise cells (see Methods). Variance reflects differences in tissue layout (domain shapes) and five random seeds used for cell assignment. Methods are ordered by the area under the ARI curve.

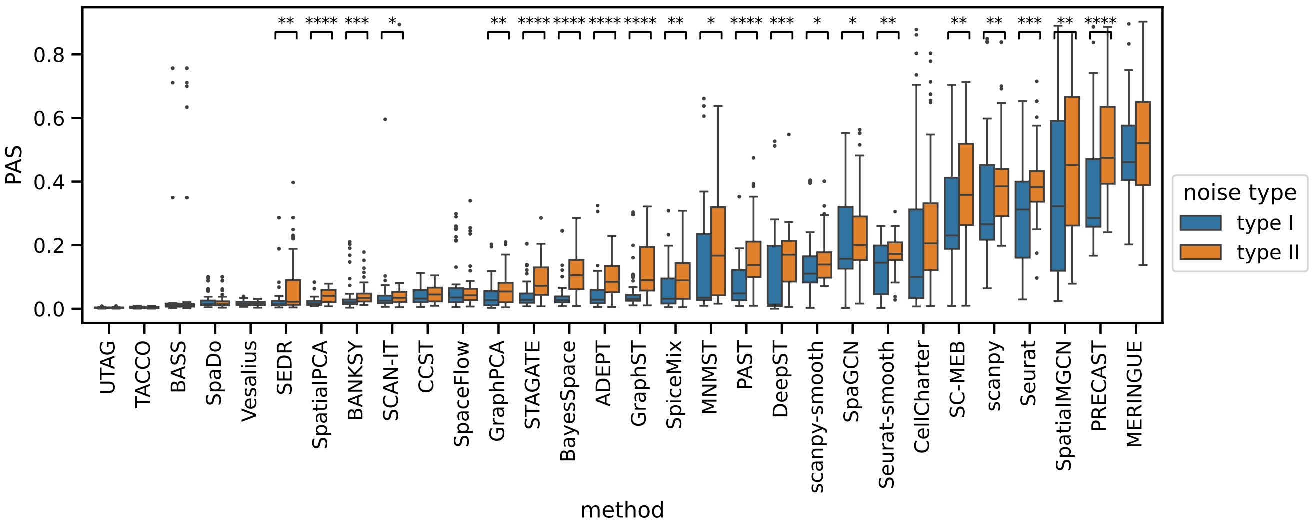


Supplementary Figure 15: PAS values across methods under types I and type II perturbations. PAS values are aggregated over all non-zero noise proportions up to, but excluding, 100%. Statistical significance was assessed using a one-sided Mann-Whitney U test.

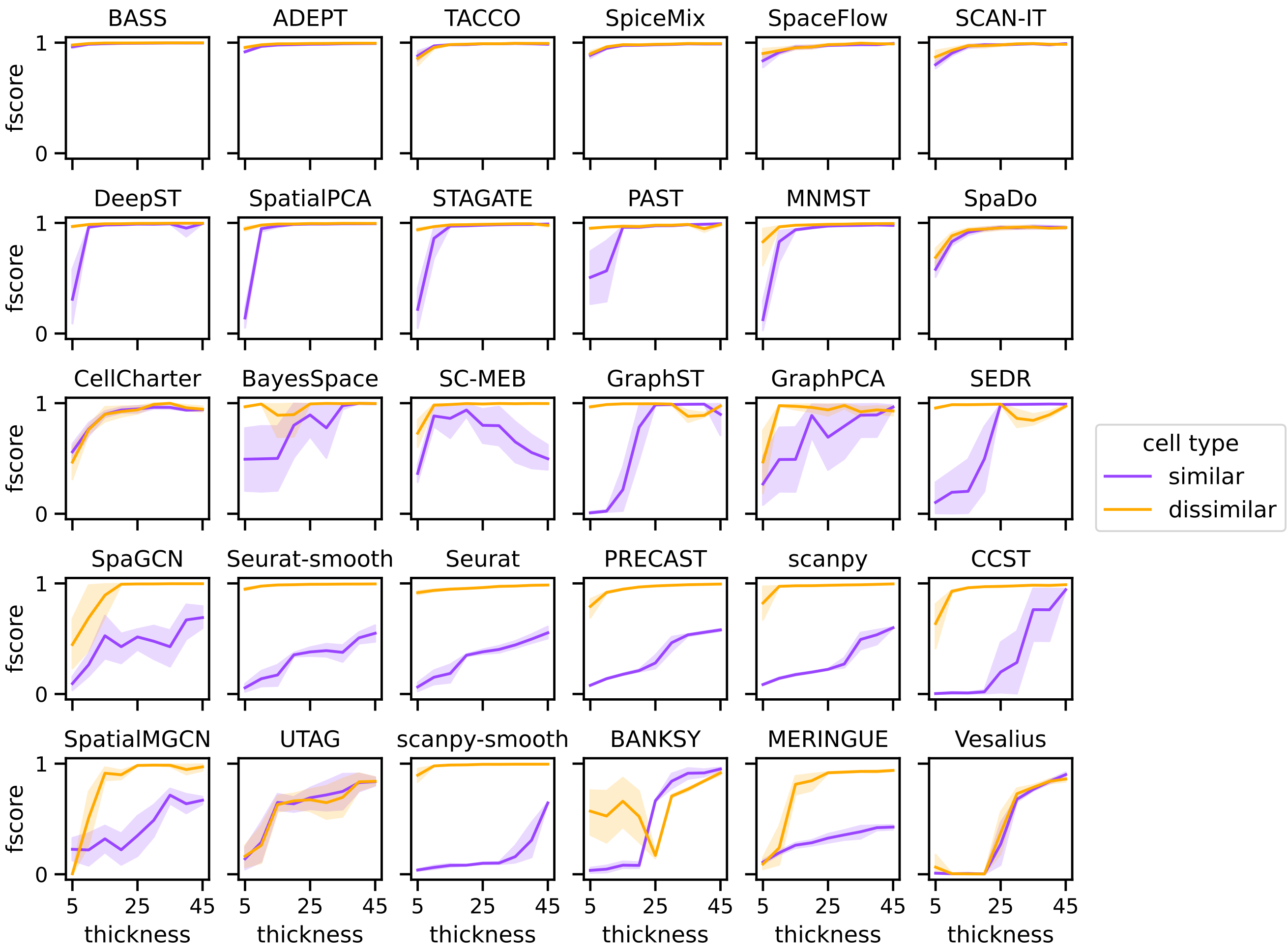


Supplementary Figure 16: Detection accuracy (F-score) across methods for the middle and right domains as a function of their thickness. The cell types assigned to these domains are either transcriptionally similar to or distinct from that of the (left-most) reference domain. Variance reflects five random seeds used for cell assignment. Methods are ordered by the area under the mean F-score curves.

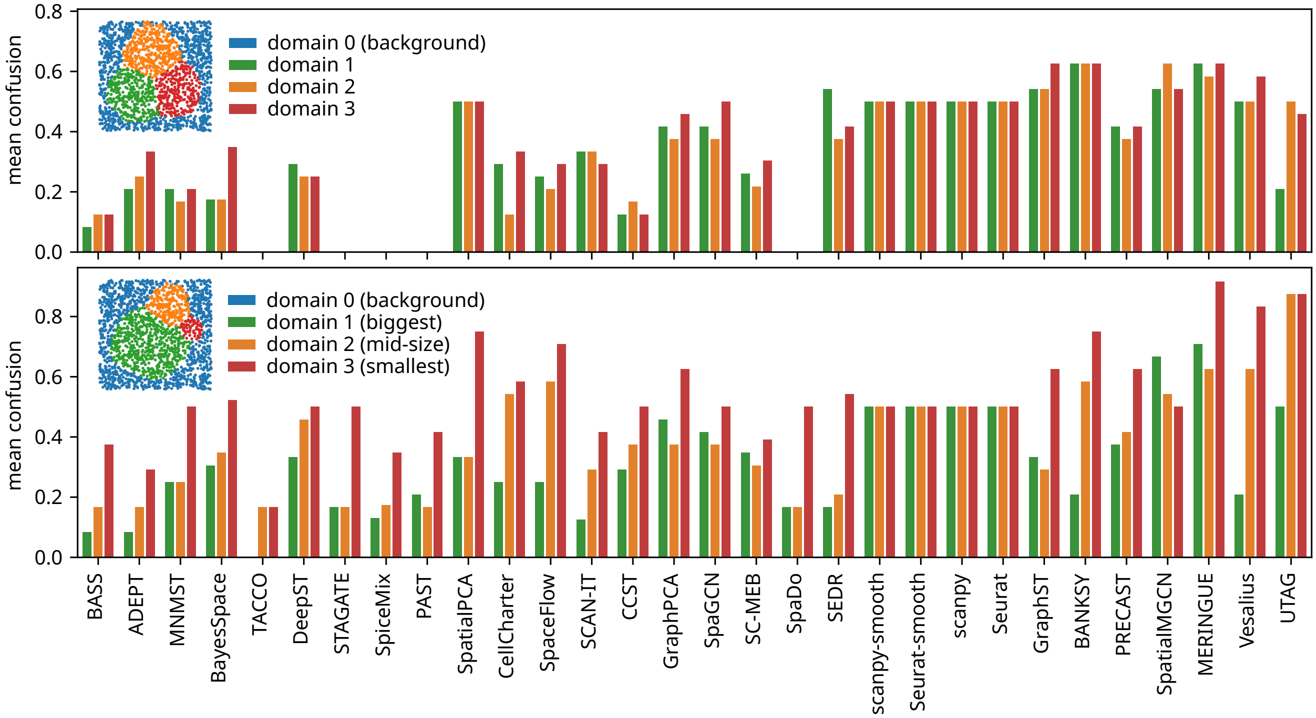


Supplementary Figure 17: Domain-wise confusion per method for equal-sized circular domains (top), and circular domains of varying size (bottom). Methods are ordered from left to right by decreasing average ARI across both layouts. Values represent the mean over all permutations of cell-type-to-domain assignments. Details of the confusion metric are provided in Methods.

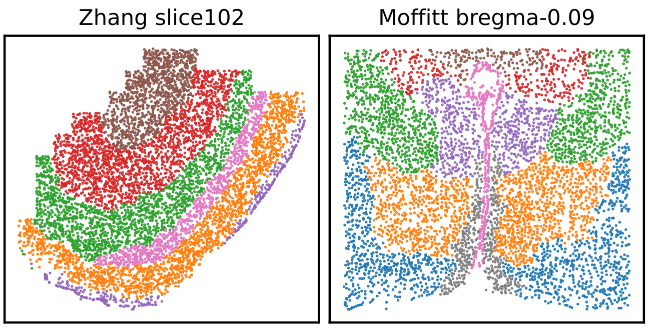


Supplementary Figure 18: Representative samples from the two MERFISH datasets illustrating differences in tissue architecture. MERFISH-Zhang exhibits a laminar layout, whereas MERFISH-Moffitt displays a more complex spatial organization.

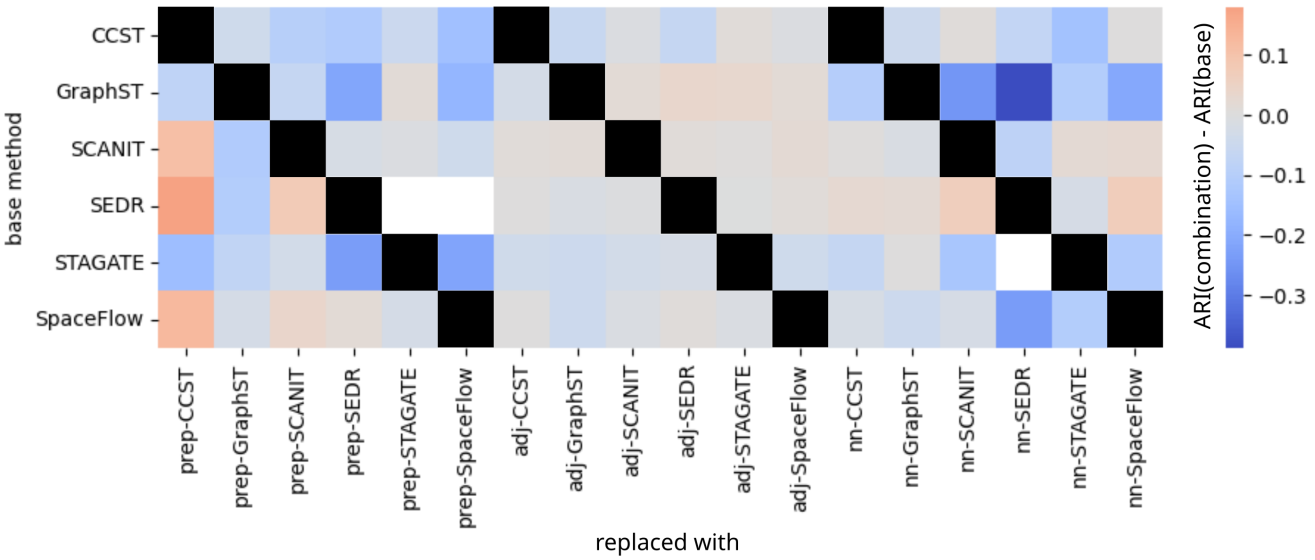


Supplementary Figure 19: Change in performance after swapping individual components in the ablation study. For each base method (rows), the preprocessing (prep), adjacency calculation and graph construction (adj), and neural-network (nn) modules were replaced with the corresponding modules from all other methods (columns). The resulting ARI values are reported as differences relative to the original method. The default clustering method is mclust for STAGATE, SEDR, and GraphST, and $k$-means for CCST, SCAN-IT, and SpaceFlow. Performance is reported as the mean ARI across the 12 Visium–Maynard samples. Blank squares indicate that no clustering output was generated.

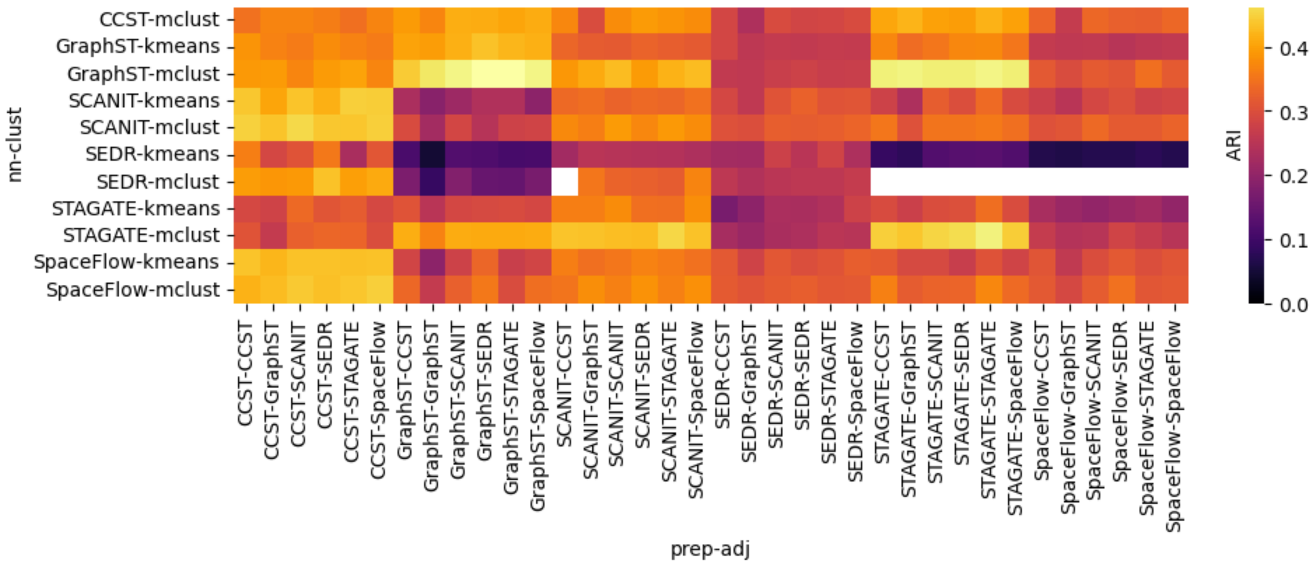


Supplementary Figure 20: Performance overview of all module combinations in the ablation study. Components for preprocessing (prep), adjacency calculation and graph construction (adj), neural network architecture and training (nn), and clustering of the learned embedding (clust) were combined across six methods, CCST, GraphST, SCAN-IT, SEDR, STAGATE, and SpaceFlow. The default clustering method is mclust for STAGATE, SEDR, and GraphST, and $k$-means for CCST, SCAN-IT, and SpaceFlow. Performance is reported as the mean ARI across the 12 Visium Maynard samples. Blank squares indicate that no clustering output was generated.

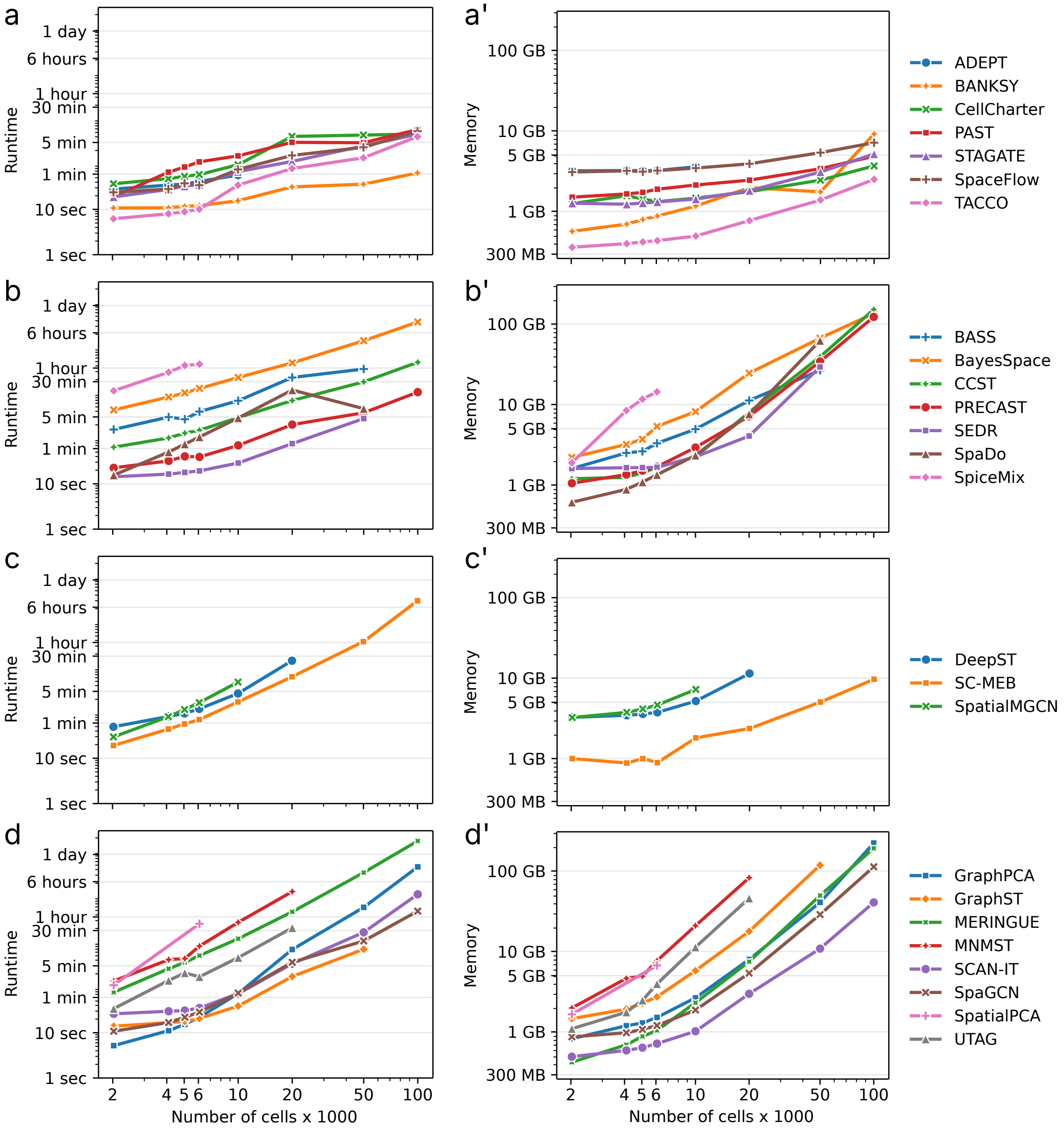


Supplementary Figure 21: Runtime (wall-clock time) and memory usage (maximum resident set size) for all methods on simulated datasets with varying numbers of cells. Unprimed panels (left) show runtime; primed panels (right) show memory usage. For visualization, methods were grouped into four categories according to their scaling trends in each quantity, using a 55/45 split for runtime and a 40/60 split for memory to obtain approximately balanced group sizes. Both axes are shown on a logarithmic scale. **a**, **a'**, slow increase in both runtime and memory. **b**, **b'**, slow increase in runtime and rapid increase in memory. **c**, **c'**, rapid increase in runtime and slow increase in memory. **d**, **d'**, rapid increase in both runtime and memory.

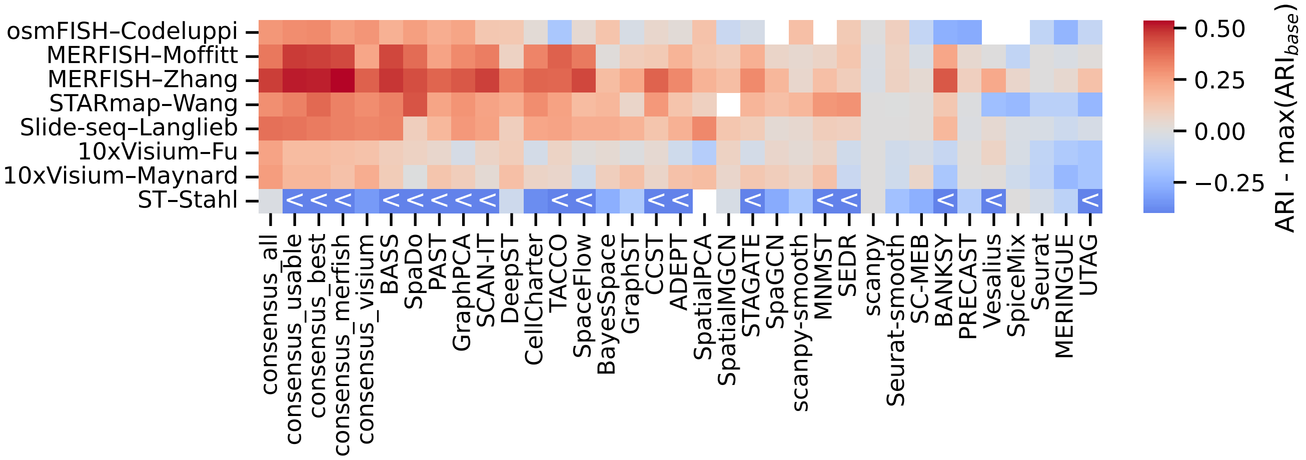


Supplementary Figure 22: Difference in ARI between each method and best-performing non-spatial baseline, as in Fig. 2a. Five additional consensus clustering results are shown: consensus over all methods, consensus over highly usable methods (PAST, CellCharter, TACCO, BASS, GraphPCA), consensus over best-performing methods (BASS, SpaDo, GraphPCA, PAST, SCAN-IT), consensus optimized for high performance on MERFISH (BANKSY, SpaceFlow, BASS, CCST, SCAN-IT) and on Visium (PAST, GraphST, SpatialPCA, MNMST, GraphPCA).

### Supplementary Tables

| Factor | Observation in real data | Result of simulation study |
| --- | --- | --- |
| Resolution | • larger improvements on high-res. datasets (Fig. 2a)<br>• method preference Visium (group V) or MERFISH (group M) by PAS (Fig. 2b)<br>• strong ARI-PAS correlation on high-res. datasets (Supplementary Fig. 5)<br>• weak ARI-PAS correlation on low-res. datasets (same Fig.) | • performance decline (group M) vs increase (group V) with decreasing resolution (Fig. 3a, Supplementary Fig. 8)<br>• subset of methods less affected (same Figs.)<br>• strong ARI-PAS correlation on high-res. datasets (Supplementary Fig. 9)<br>• weak ARI-PAS correlation on low-res. datasets (same Fig.) |
| Panel size | • smaller gains on osmFISH than other high-res. datasets (Fig. 2a) | • group M with more moderate performance decline with shrinking panel size (Fig. 3b, Supplementary Fig. 8)<br>• sharp drop in performance for BANKSY and CellCharter (same Fig.) |
| Sparsity | • low accuracy on Slide-seq datasets (Supplementary Fig. 3) | • highly sensitive vs more robust subsets of methods (Supplementary Fig. 10)<br>• most of sparsity-resistant methods perform better on Slide-seq (Fig. 2a, Supplementary Figs. 3 and 10) |
| Domain size | • missed L4 domain in Visium-Maynard dataset (Supplementary Fig. 6) | • 6 characteristic behaviors dependent on domain thickness (Figs. 4a,b)<br>• dependence modulated by transcriptional distinctness (Supplementary Fig. 16 |
| Distinctness | • missed L4 domain in Visium-Maynard dataset (Supplementary Fig. 6) | • 3 failure modes with increasing transcriptional similarity (Fig. 3c)<br>• failure modes linked to imposed spatial coherence (same Fig.)<br>• transcriptional heterogeneity impacts performance more than similarity (Supplementary Figs. 11 and 14)<br>• small subsets of robust methods in both cases (same Figs.) |
| Domain shape | • higher accuracy on laminar brain layers (Supplementary Figs. 3 and 18) | • Most methods prefer laminar shapes (Fig. 4c)<br>• UTAG performs competitively only on laminar shapes (same Fig.) |

Supplementary Table 1: Summary of performance dependencies observed in real spatial transcriptomics data and their systematic validation using semi-synthetic simulations.

| Method | Preprocessing | Neighborhood & Graph | Neural network & training |
|---|---|---|---|
| **STAGATE** | • Top 3000 HVGs<br>• Log-transform<br>• Normalize library size to $10^4$ | • Top 6-NN | • Autoencoder with edges between 6-NN cells<br>• Minimize reconstruction loss<br>• Return hidden embedding |
| **SCAN-IT** | • Normalize library size to 1<br>• Fit SOM; select 3000 most variable genes from SOM<br>• Log-transform; standardize (zero mean, unit variance) | • Alpha-complex spatial graph<br>• Top 5-NN | • Deep Graph Infomax (maximize MI)<br>• Negatives: perturb gene expressions<br>• Postprocess with metric MDS to 30 dimensions |
| **SEDR** | • Exclude genes: detected in $< 50$ cells or $< 10$ total counts<br>• Normalize library size to $10^6$<br>• Top 2000 HVGs (`seurat_v3`)<br>• 200 PCs | • Top 12-NN | • VGAE<br>• Jointly embed adjacency and gene expressions<br>• Concatenate, then refine with DEC (SpaGCN-style clustering) |
| **CCST** | • Exclude genes: detected in $< 3$ cells<br>• Normalize library size to 1<br>• Standardize (zero mean, unit variance)<br>• 200 PCs | • Radius such that avg number of neighbors $\approx 3$<br>• Let $A$ be adjacency; use $\tilde{A} = (1 - \lambda)A + \lambda I$ | • Deep Graph Infomax (maximize MI)<br>• Negatives: perturb $\tilde{A}$ edges<br>• After embedding: PCA to 30 dimensions |
| **GraphST** | • Top 3000 HVGs (`seurat_v3`)<br>• Normalize library size to $10^4$<br>• Log-transform<br>• Scale (no zero-centering; cap at 10) | • Top 3-NN<br>• Symmetrize adjacency | • Autoencoder-like Deep Graph Infomax (maximize MI) + reconstruction loss (minimize MSE)<br>• After embedding: PCA of reconstructed expression to 30 dimensions |
| **SpaceFlow** | • Exclude genes with $< 1$ count<br>• Normalize library size to $10^4$; Log-transform<br>• Keep top 3000 HVGs (`cell_ranger`; subset)<br>• 50 PCs | • Alpha-complex spatial graph; remove self-edges<br>• Uses Top 10-NN distances to set alpha scale (heuristic) | • Deep Graph Infomax (maximize MI) + spatial regularization parameter (regularizer)<br>• Embedding dimension 50 |

Supplementary Table 2: 6 neural-network-based methods (rows) were decomposed into 3 components (columns). HVG, highly variable genes; SOM, self-organizing map; MI, mutual information; MDS, multi-dimensional scaling; PCs, principal components; VGAE, variational graph autoencoder; DEC, Deep Embedded Clustering.

| Method | Availability | | | Maintenance | | | Dokumentation | | |
|---|---|---|---|---|---|---|---|---|---|
| | 1(a) | 1(b) | 1(c) | 2(a) | 2(b) | 2(c) | 3(a) | 3(b) | 3(c) |
| ADEPT | 0 | 1 | 1 | 0 | 1 | 0 | 1 | 0 | 1 |
| BASS | 0 | 0 | 1 | 1 | 1 | 1 | 1 | 0 | 1 |
| BANKSY | 0 | 1 | 1 | 1 | 1 | 0 | 0 | 0 | 1 |
| BayesSpace | 0 | 1 | 1 | 1 | 0.5 | 1 | 1 | 1 | 1 |
| CCST | 0 | 1 | 0 | 0 | 0.5 | 1 | 0 | 0 | 0.5 |
| CellCharter | 1 | 0 | 1 | 1 | 1 | 1 | 1 | 1 | 1 |
| DeepST | 1 | 1 | 1 | 1 | 0 | 0 | 0 | 0 | 0.5 |
| GraphPCA | 1 | 0.5 | 0.5 | 1 | 0.5 | 1 | 1 | 0 | 1 |
| GraphST | 1 | 1 | 1 | 1 | 0.5 | 1 | 1 | 0 | 1 |
| MERINGUE | 0 | 0 | 1 | 1 | 1 | 0 | 0 | 0 | 1 |
| MNMST | 1 | 1 | 1 | 1 | 1 | 1 | 0 | 0 | 1 |
| PAST | 1 | 0.5 | 1 | 1 | 1 | 1 | 1 | 1 | 1 |
| PRECAST | 1 | 0.5 | 1 | 1 | 1 | 1 | 1 | 1 | 1 |
| SCAN-IT | 0 | 0 | 0 | 1 | | 0 | 0 | 0 | 1 |
| SC-MEB | 1 | 0.5 | 1 | 1 | 1 | 0 | 1 | 1 | 1 |
| SEDR | 0 | 1 | 1 | 1 | 1 | 0 | 1 | 0 | 1 |
| SpaGCN | 1 | 1 | 1 | 1 | 1 | 1 | 0 | 0 | 1 |
| SpaceFlow | 1 | 1 | 1 | 1 | 0.5 | 1 | 0 | 0 | 1 |
| SpaDo | 0 | 0 | 1 | 0 | 1 | 1 | 0 | 0 | 1 |
| SpatialMGCN | 0 | 1 | 0 | 1 | | 0 | 0 | 0 | 0.5 |
| SpatialPCA | 0 | 0 | 1 | 1 | 1 | 1 | 1 | 1 | 1 |
| SpiceMix | 0 | 0.5 | 0.5 | 1 | 0.5 | 1 | 0 | 0 | 0.5 |
| STAGATE | 0 | 1 | 0.5 | 1 | 0.5 | 1 | 1 | 0 | 1 |
| TACCO | 1 | 1 | 1 | 1 | 1 | 1 | 1 | 1 | 1 |
| UTAG | 1 | 1 | 1 | 1 | 1 | 1 | 0 | 1 | 1 |
| Vesalius | 0 | 0.5 | 1 | 1 | 1 | 1 | 1 | 1 | 1 |

Supplementary Table 3: All benchmarked methods are open-source and available on GitHub. Usability was evaluated according to three criteria: 1. Availability: (a) installable via a package manager; (b) dependencies clearly listed; (c) installation instructions provided. 2. Maintenance: (a) modular code structure; (b) active response to issues on GitHub; (c) version control in use. 3. Documentation: (a) dedicated package documentation (e.g., via readthedocs.io); (b) well-documented functions; (c) tutorials illustrating how to run the method.

### Supplementary Note 1: A conceptual overview and taxonomy of neural network–based domain detection methods

**Preprocessing** With the exception of DeepST, all methods follow a standard preprocessing workflow that includes filtering genes and cells with low counts, normalization, and log-transformation of gene expression. DeepST deviates from this pattern by skipping these steps and instead scaling the gene counts of each cell according to its correlation with its nearest neighbors.

After preprocessing, SpaGCN and CCST compute principal components (PCs), whereas other methods, including SCAN-IT, STAGATE, SpaceFlow, ADEPT, PAST, SpatialMGCN, GraphST, SEDR and MNMST, select highly variable genes (HVGs). Several methods then apply additional transformation. SEDR performs PCA on the HVGs; SCAN-IT applies self-organizing maps (SOMs)[1] to identify spatially variable genes; and MNMST enhances spot expression values using BANKSY.

**Neighborhood graph construction** Except for SpaGCN, which constructs a complete graph, all neural-network–based methods build a sparse neighborhood graph using some variant of nearest-neighbor selection. SpaceFlow and SCAN-IT construct an alpha-complex graph[2], ADEPT, STAGATE, Spatial-MGCN, and CCST define neighbors using a fixed spatial radius, and DeepST, SEDR, PAST, GraphST, and MNMST rely on $k$-nearest neighbors ($k$NN). Several methods additionally weight graph edges based on spatial distance to better preserve spatial proximity (SpaGCN, PAST, and MNMST). CCST further introduces a trade-off parameter that controls the relative influence of spatial versus gene expression similarity, enabling adaptive integration of the two information sources.

**Architecture and Training** We categorize neural-network–based methods according to their core architectural paradigms and corresponding training strategies:

1. *Autoencoder-based methods*

   Autoencoder approaches learn low-dimensional latent representations by compressing and reconstructing the input data through an encoder–decoder architecture.

   STAGATE and ADEPT fall into this category. Both employ graph autoencoder architectures with attention mechanisms to model similarity between neighboring spots, and optimize reconstruction of gene expression data using a mean-squared-error loss function.

   In addition, ADEPT identifies differentially expressed genes and iteratively refines both gene selection and clustering to reduce within-cluster variance.

2. *Variational graph autoencoder (VGAE) methods*

   Variational graph autoencoders extend autoencoders to graph-structured data using variational inference[3] to learn node embeddings that capture uncertainty and probabilistic relationships.

   DeepST, SEDR, PAST, and Spatial-MGCN, all integrate VGAE-style components into an autoencoder framework. SEDR incorporates a masked vector into the architecture, and PAST augments the model with a Bayesian neural network coupled with a self-attention mechanism.

   In addition to the conventional mean-squared-error loss used for the autoencoder component, SEDR, DeepST, and PAST apply the Kullback-Leibler divergence penalty assuming a standard normal prior over the latent variables. In contrast, Spatial-MGCN models expression counts using a zero-inflated negative binomial (ZINB) distribution to better account for zero inflation and dropout in spatial transcriptomics.

   PAST and Spatial-MGCN introduce additional regularization terms that encourage separation of distinct domains by preserving pair-wise distances in the embedding space. Furthermore, PAST employs ripple walk sampling[4] to enable scalable training on large graphs.

3. *Deep Graph Infomax (DGI)-based methods*

---

[1] Self-organizing maps are unsupervised neural networks that project high-dimensional data to a lower-dimensional grid while preserving topological structure.

[2] An alpha-complex graph is a subgraph of the Delaunay triangulation that captures the shape and connectivity of a point set.

[3] Variational inference approximates the posterior distribution over latent variables by optimizing a tractable variational distribution.

[4] Ripple walk sampling is a random walk-based method for graph sampling that explores nodes by iteratively expanding a "rippl" from a starting node, visiting neighboring nodes at increasing distances in a controlled manner.

Deep Graph Infomax (DGI) [3] is a self-supervised graph representation learning paradigm that maximizes the mutual information between global graph summaries and local node embeddings. Negative samples are constructed by corrupting the graph structure or features, and the model uses a graph neural network (GNN) encoder to learn embeddings that discriminate real from corrupted graph contexts.

CCST, GraphST, SCAN-IT, and SpaceFlow belong to this class but differ in corruption strategies, loss design, and postprocessing.

- CCST perturbs the graph by permuting edges rather than node attributes to generate negative samples.
- SpaceFlow adds a spatial regularization term that penalizes embedding similarity between spatially distant cells.
- GraphST integrates DGI within an autoencoder, combining three losses: reconstruction loss on the decoder, a DGI loss on the encoder, and a symmetric contrastive loss.
- CCST and SCAN-IT both apply standard DGI losses and further project embeddings through PCA (CCST) or metric multidimensional scaling (SCAN-IT) to enhance cluster stability.

4. *Pure graph convolutional network (GCN) method*

   SpaGCN is the only method in this benchmark that uses a pure GCN framework. It integrates gene expression, spatial location, and histological information within a GCN and iteratively optimizes cluster separation using Deep Embedded Clustering (DEC).

5. *Non-negative matrix factorization (NMF)-based method*

   MNMST jointly factorizes spatial and expression graphs using a non-negative matrix factorization multi-layer network, learning shared low-dimensional features while incorporating sparse self-representation learning to infer expression adjacency.

### Supplementary Note 2: Details of real datasets and domain annotations

This section provides details of the real datasets included in the benchmark, including the origins of the domain annotations used as ground truth for evaluation.

**osmFISH Codeluppi** The osmFISH dataset of the mouse somatosensory cortex was downloaded from linnarssonlab.org/osmFISH/availability in loom format. The count matrix and cell coordinates were extracted and saved as csv files. Domain annotations are provided with the dataset and are based on marker genes [4].

**MERFISH Moffitt** The MERFISH dataset of the mouse hypothalamic preoptic region was downloaded from datadryad.org/dataset/doi:10.5061/dryad.8t8s248 in csv format [5]. Counts and cell coordinates for each sample were saved separately as csv files. Five samples were annotated by Li *et al.* and their domain annotations are available at github.com/zhengli09/BASS-Analysis/tree/master/data [6]. Annotations are based on spatial gene expression patterns and histology, with reference to the Allen Brain Atlas [7].

**MERFISH Zhang** The MERFISH dataset of the mouse primary motor cortex is available via doi.brainimagelibrary.org/doi/10.35077/g.21 and was downloaded from download.brainimagelibrary.org/cf/1c/cf1c1a431ef8d021 in h5ad format [8]. Counts and cell coordinates for each sample were extracted and saved separately as csv files. 33 samples were annotated by Xu *et al.* based on marker genes, and these domain annotations are available at zenodo.org/records/8316334 in h5ad format [9].

**STARmap Wang** The STARmap dataset of the mouse primary visual cortex was downloaded from zenodo.org/records/10698912 in h5ad format. The count matrix and cell coordinates were extracted and saved as csv files. Domain annotations are provided with the dataset and are based on marker genes and anatomical annotation [10].

**Slide-seq Langlieb** The Slide-seq dataset of the whole mouse brain is available via braincelldata.org and was downloaded from drive.google.com/drive/folders/1-3eLeLQxg2K2JEgxV_7eUg5JgDB6N7tg in h5ad format [2]. Counts and bead coordinates for each sample were extracted and saved separately as csv files. Domain annotations are provided with the dataset and are based on label transfer from the Allen Brain Atlas and marker genes [7]. In this study, we used the first 10 pucks (01-10). In addition, only beads assigned to the left hemisphere (indicated by `CCF_LeftRight`) were included in the analysis.

**Visium Fu** The Visium dataset of human breast cancer was downloaded from support.10xgenomics.com/spatial-gene-expression/datasets/1.1.0/V1_Breast_Cancer_Block_A_Section_1 in HDF5 format. The count matrix and spot coordinates were extracted and saved as csv files. The sample was annotated by Xu *et al.* based on pathological features and cell-type annotations, and the domain labels are available at github.com/JinmiaoChenLab/SEDR_analyses/blob/master/data/BRCA1/metadata.tsv [11].

**Visium Maynard** The Visium dataset of the human dorsolateral prefrontal cortex was downloaded from research.libd.org/spatialLIBD in HDF5 format for each sample [1]. Count matrices and spot coordinates were extracted and saved as csv files. Domain annotations are provided with the dataset and are based on histology, t-SNE visualization of domain-specific marker genes, and marker gene expression patterns.

**ST Stahl** The Spatial Transcriptomics (ST) dataset of the mouse olfactory bulb was downloaded from spatialresearch.org/resources-published-datasets/doi-10-1126science-aaf2403 in tsv format.

### Supplementary Note 3: Implementation inconsistencies and reproducibility considerations

Across multiple methods in this benchmark, we observed reproducibility-relevant inconsistencies between the mathematical descriptions provided in the associated manuscripts and the corresponding publicly released implementations. The list below did not arise from a systematic investigation and is therefore likely incomplete.

For SpaGCN, the published formulation of the graph convolutional network (GCN) layer and the parameterization of the Student's t-distribution differ from the implementation. For example, the implemented GCN layer omits the non-linearity described in the manuscript, and the Student's t-distribution uses a hard-coded value of $\alpha$ that does not match the reported definition.

Similarly, CCST stacks multiple graph convolutional operations without intermediate non-linear activations. This renders the effective transformation equivalent to a single linear layer, which is not apparent from the textual description.

Both STAGATE and CCST construct spatial graphs using radius-based neighborhood definition graphs with dataset-specific or, in the case of CCST, hardcoded distance thresholds. Because absolute spatial distances are not standardized across technologies, selecting an appropriate radius is non-trivial and can materially influence results, whereas $k$-nearest-neighbor-based graphs may provide greater robustness across datasets.

Several methods also contain hard-coded internal parameter values that are undocumented, limiting the user's ability to tune models and potentially posing challenges for less-experienced users attempting to apply the tools in new settings.

Finally, for GraphST, the manuscript alternates between descriptions of augmentation-based and augmentation-free contrastive learning, and key preprocessing steps (such as dimensionality reduction of reconstructed expression data to a fixed low rank using PCA) are either underspecified or unconventional.
